# Molecular and spatial design of early skin development

**DOI:** 10.1101/2022.12.28.522081

**Authors:** Tina Jacob, Karl Annusver, Paulo Czarnewski, Tim Dalessandri, Maria Eleni Kastriti, Chiara Levra Levron, Marja L Mikkola, Michael Rendl, Beate M Lichtenberger, Giacomo Donati, Åsa Björklund, Maria Kasper

## Abstract

A wealth of specialized cell populations within the skin facilitates its hair producing, protective, sensory and thermoregulatory functions, but how the vast cell-type diversity and tissue architecture develops is largely unexplored. Here, with single-cell transcriptomics, spatial cell-type assignment and cell-lineage tracing we deconstruct early embryonic mouse skin during the key transitions from seemingly uniform developmental precursor states to a multilayered, multilineage epithelium and complex dermal identity. We reveal the spatiotemporal emergence of hair-follicle-inducing, muscle-supportive, and fascia-forming fibroblasts. We also uncover the formation of the panniculus carnosus muscle, sprouting blood vessels without pericyte coverage, and the earliest residence of mast and dendritic immune cells in skin. Finally, we reveal an unexpected epithelial heterogeneity within the early single-layered epidermis and a signaling-rich periderm layer. Overall, this cellular blueprint of early skin development establishes histological landmarks – essential for placing cells in their spatial tissue context – and highlights unprecedented dynamical interactions among skin cells.

## INTRODUCTION

During skin development, one of the most remarkable changes occurs when the epidermis transforms from a single-layered epithelium, to a multi-layered and appendage-producing epithelium. Mouse epidermis develops from the surface ectoderm at embryonic day (E) 9.5, starting as a single layer of basal keratinocytes that is subsequently covered by a transient protective layer of squamous cells called ‘periderm’. Within the following 10 days (i.e. until birth at approx. E19.5) a fully stratified epidermis is formed, which acts as a reliable barrier keeping pathogens outside and water inside (De Falco et al., 2014). During these 10 days, also the epidermal appendages form. In dorsal skin, hair follicles develop in three waves, with the first epithelial thickenings – so-called hair placodes – being morphologically visible at E14.5. Hair placodes maintain a tight dialogue with the underlying dermal condensate (DC), a mesenchymal signaling center that stays in close contact with the hair follicle throughout its lifetime (Biggs and Mikkola, 2014; Saxena et al., 2019; Schmidt-Ullrich and Paus, 2005; Sennett and Rendl, 2012). The vast majority of studies on embryonic skin to date have focused on the skin’s epidermis (Fuchs, 2007; Hu et al., 2018; Sotiropoulou and Blanpain, 2012). Nevertheless, important aspects of epidermal development remain unresolved, such as the maturation of the periderm and its signaling potential, basal-cell heterogeneity prior to placode formation, and the involvement of mature placode cells in shaping the skin’s dermal architecture.

In the dermis, fibroblasts are the most abundant cell type, yet little is known about their heterogeneity and contributions to early skin development. In recent years, fibroblasts have been increasingly recognized as important modulators of epithelial stem cells and their niches (Plikus et al., 2021). From studies in adult organs such as the skin, intestine, heart and mammary gland, we now know that fibroblasts play an important role in tissue homeostasis, wound repair as well as cancer formation (David et al., 2020; Furtado et al., 2016; Griffin et al., 2020; Houthuijzen and Jonkers, 2018; Kobayashi et al., 2019; Nelson and Bissell, 2006; Rognoni and Watt, 2018; Roulis and Flavell, 2016; Sasaki et al., 2018; Souders et al., 2009; Sumbal and Koledova, 2019). However, the diverse functional roles that fibroblasts play during organ development and their importance are largely unexplored.

The few studies that focused on the developing dermis were mostly centered on the molecular and cellular establishment of hair follicles (e.g., Biggs et al., 2018; Ge et al., 2020; Gupta et al., 2019; Mok et al., 2019) leaving a major gap in knowledge about the non-hair-follicle-related mesenchymal cell types during early skin development. Based on antibody staining, *in vivo* lineage-tracing and skin reconstitution experiments, it has been proposed that dermal fibroblasts derive from a single fibroblast lineage that diverges at E16.5; one forming the upper (papillary) dermis including the hair follicle-associated dermal papillae, dermal sheath and arrector pili muscles, and one giving rise to the lower (reticular) dermis and adipocytes of the hypodermis (Driskell et al., 2013). Although the existence of fibroblast heterogeneity and potential fate-specification prior to the lineage divergence at E16.5 has been proposed (Rinkevich et al., 2015), major questions remain. How heterogeneous are fibroblasts during early skin development? When does fibroblast heterogeneity emerge? By which means do early fibroblasts support tissue specification and maturation? Importantly, with this work we did not only study the skin but also exploited it as a prime organ to explore the variety and functions of fibroblasts in embryonic organ development.

A major challenge to answer any of these questions is the complete lack of histological or molecular tissue landmarks in early developing skin. In adult mouse skin, the only certain landmark to date that defines “skin-associated” cells is the panniculus carnosus muscle (PCM). Only the cells above the PCM (epidermis/dermis), the PCM itself, and a thin layer of connective tissue cells (called fascia) just below the PCM are considered “skin-associated” cells. Surprisingly, when the PCM is formed has not been reported. At E12.5, the future skin dermis and fascia, as well as non-skin-associated cells, are part of a seemingly homogenous tissue space spanning from the vertebrae to the epidermis. Similarly, at E13.5 and 14.5, little is known about dermal tissue architecture and cell type diversity, asking for more answers. When does the PCM form? What is the spatiotemporal diversity of all other cell types, such as neural crest-derived, vessel-associated, or immune cells, during early skin development?

This work aims at presenting a holistic picture of early skin development. Through comprehensive computational analysis of all cell types sampled at E12.5, E13.5, E14.5, cell-type localization *in situ* and *in vivo* cell-fate mapping, we i) determined fibroblast heterogeneity and onset of lineage commitment, ii) resolved the periderm-transcriptome and epidermal cell heterogeneity prior to placode formation, iii) characterized all other major cell types, iv) portrayed the comprehensive interplay between skin cell types, and v) provided new histological landmarks which are essential to place cells in their spatial tissue context.

## RESULTS

### Single-cell profiling-assisted generation of histological landmarks in E12.5, E13.5 and E14.5 skin

To determine the cellular composition of early embryonic skin, and to unveil decisive signaling events driving early skin maturation, we profiled E12.5, E13.5 and E14.5 mouse back skin. We isolated full-thickness dorsal skin and generated single-cell transcriptome libraries of epithelial and stromal cells using the 10x (v2) Genomics Chromium system. To assure true biological replicates, 5 embryos per embryonic time point were processed, sequenced, and quality-controlled individually (**Figure S1A-B**). We also embedded the remaining body of each sequenced embryo as well as intact littermates for histology to ensure that the sequenced embryos had been developed as expected for their respective age (**Figure S1C**). After quality control, all three timepoints were analyzed together (**Figure S1D-F**; **Methods**) in order to better capture developmental trends and the dynamics of disappearing and newly emerging cell populations.

The complete dataset contains 32,194 single-cell transcriptomes with 11,280 cells coming from E12.5, 9,964 cells from E13.5 and 10,950 cells from E14.5 (**Figure 1A**). Based on cluster-specific gene expression we identified keratinocytes, fibroblasts, immune cells, vessel-associated mural (pericytes and vascular smooth muscle cells) and endothelial cells, neural crest-derived melanocytes and Schwann cells, and muscle cells (**Figure 1B-C**; **Table S1**). Through fluorescent *in situ* hybridization (FISH for mRNA) and immunofluorescence staining (IF for protein) of cell-type specific marker genes, we mapped all major cell types within E12.5, E13.5, and E14.5 skin tissue (**Figure 1D-L**). We also used these cell-type-specific markers together with wheat germ agglutinin (WGA) cell-membrane staining to establish histological landmarks of early skin development, which the commonly used H&E staining cannot resolve (**Figure S1G-I**). Notably, staining for ACTC1 revealed the stepwise development of multiple muscle layers including the PCM (the latter is only evident from E14.5) (**Figure 1D-F**), PTPRC highlights the exclusively dermal location of immune cells (**Figure 1J**), *Sox10* shows large nerve trunks growing towards the epidermis at E12.5 and more spread-out nerves at E14.5 (**Figure 1K-L**), and *Rgs5/Pecam1* co-staining depicts the dense vascular network (**Figure 1G-I**). For this work, these newly established landmarks (summarized in **Figure 1M-O**) were instrumental for the correct mapping and placement of numerous cell populations within the rapidly developing full-thickness skin, which we present in the following sections.

**Figure 1.**
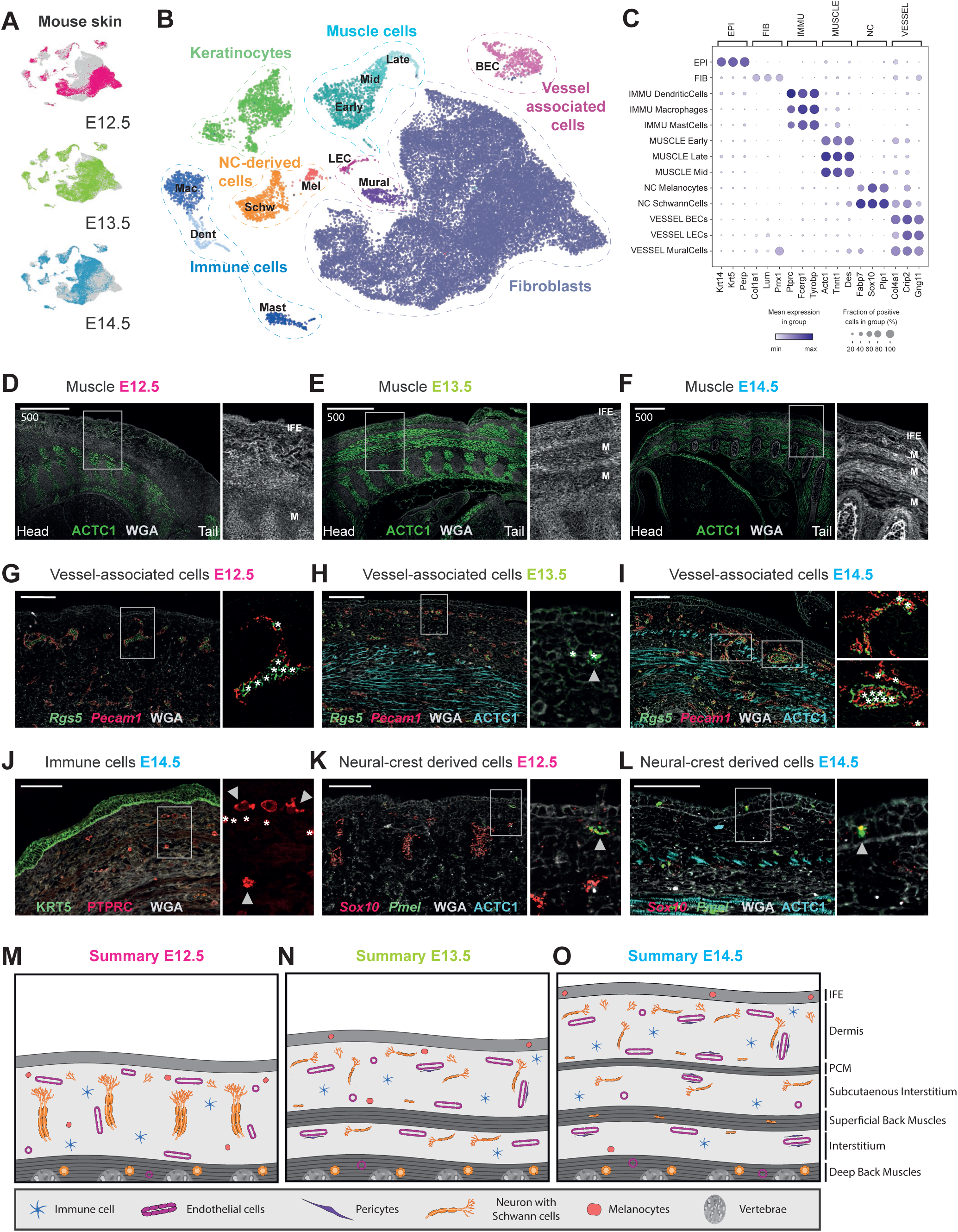
Anatomy of embryonic skin from E12.5, E13.5, and E14.5. (A) UMAP visualization of all cells. Highlighted in color are the cells from the different embryonic ages (n = 11 280 cells for E12.5, 9 964 for E13.5, and 10 950 for E14.5). (B) UMAP visualization of all cells, colored according to first-level clustering (n = 32 194 cells). Fibroblast subclusters and keratinocyte subclusters were merged and analysed in detail later (Figure 2, 3 and 6, respectively). (C) Dot plot depicting expression of unifying markers for main cell classes. (D-L) mRNA and protein stainings (RNA FISH in italics; protein IF in capital letters) revealing the major anatomical landmarks of dorsal embryonic skin. Shown are dorsal areas of full embryo FFPE sagital sections. Microscope images originate from larger tile scans (n = 3 mice). Scale bars, 500μm (D-F) and 100μm (G-L). (D-F) ACTC1 stains muscle layers. Zoom in to visualize how membranous WGA staining reveals anatomical layers of embryonic skin. M marks developing muscle layers. (G-I) *Pecam1* stains endothelial cells and *Rgs5* stains mural cells. Zoom in on single vessels to visualize emergence of mural cells. Note that intravascular erythrocytes show strong autofluorescence (asterisks in G-I). Arrowhead in (H) marks earliest evidence of mural cells at E13.5. Upper zoom in in (I) shows smaller vessel with discontinuous mural cell lining while lower zoom in in (I) shows larger vessel with continuous mural cell lining. (J) PTPRC stains immune cells and KRT5 stains epidermis. Zoom in to highlight immune cells with dendritic phenotype (arrowhead). Note that intravascular erythrocytes show strong autofluorescence in green and red (in zoom in highlighted with asterisks). (K-L) *Sox10* stains melanocytes and Schwann cells and *Pmel* stains melanocytes only. Zoom in to showcase the arrival of melanocytes in the epidermis (arrowhead). (M-O) Schemes summarizing anatomical landmarks at E12.5, E13.5 and E14.5, respectively.

### Fibroblast heterogeneity exists long before the reported establishment of papillary and reticular dermis

We focused our initial analysis on the most abundant cell type in developing skin – the fibroblasts – as it was highly conceivable that they are actively involved in orchestrating the stepwise maturation of skin to its full function, such as the establishment of vessels, fat, and epithelial appendages. Our dataset contained in total 25,944 fibroblasts out of 32,194 randomly sampled cells. The contribution of high-quality cells from 15 individual embryos allowed for robust identification of 22 fibroblast subpopulations (**Figure S2A**; **Table S2**). Overall, fibroblasts isolated from the skin and underlying (non-skin) tissue (**Methods** and **Discussion**) separated into seven major groups of cell populations. We named these groups *FIB Origin, FIB Deep, FIB Upper & DC, FIB Lower, FIB Muscle, FIB Inter* and *CHOND* (**Figure 2B, 3B** and **S2A**) based on several criteria such as their appearance in development (**Figure 2A**), their gene expression profiles (**Figure 2C** and **3C; Table S1**), RNAvelocity analysis (**Figure S2B**), and tissue location, all of which are discussed in more detail in the following paragraphs.

**Figure 2.**
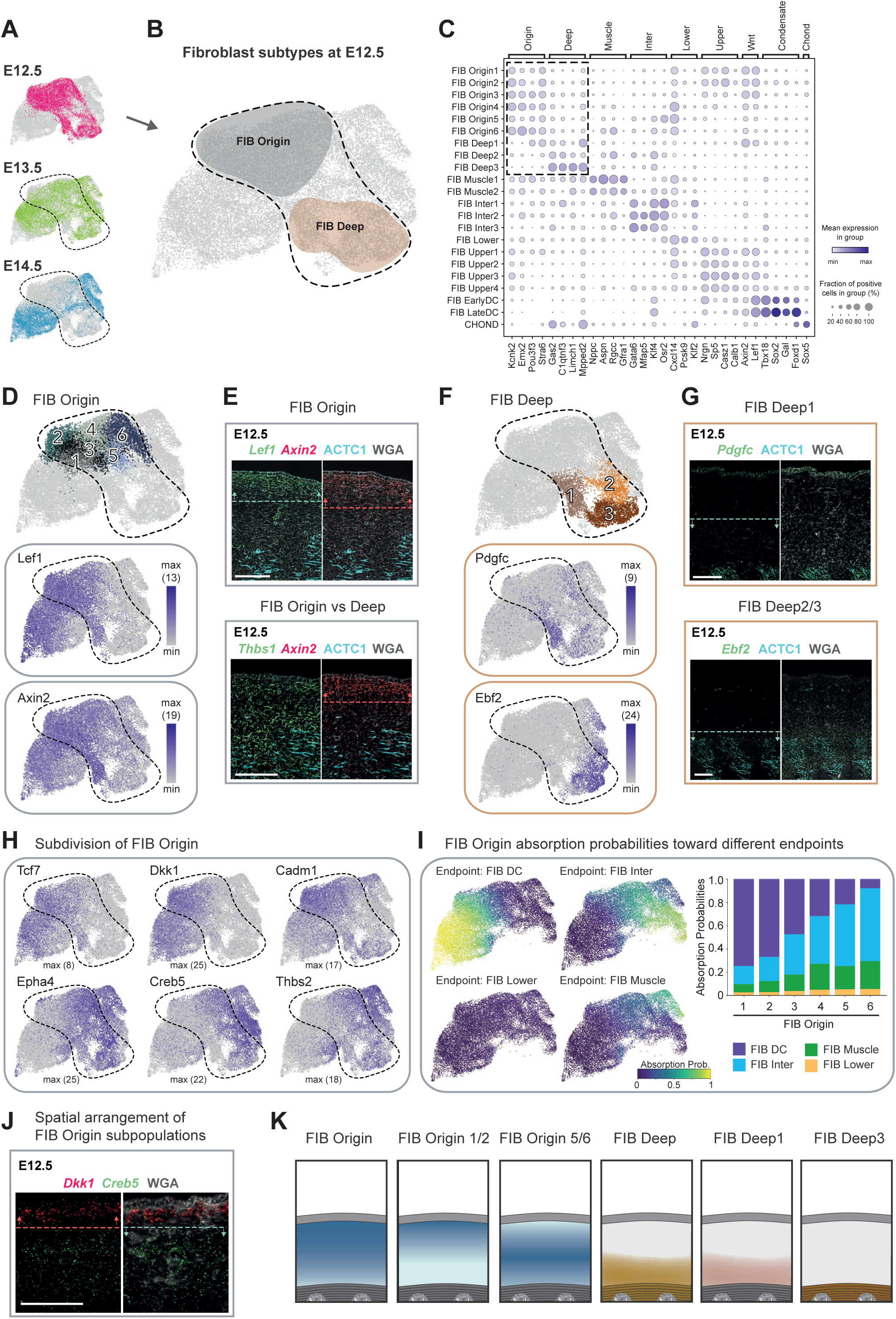
Deconstruction of fibroblast heterogeneity at E12.5 (expression and location) (A) UMAP visualization of all fibroblasts. Highlighted in color are the cells from the different embryonic ages (n = 10 008 cells for E12.5, 8 016 for E13.5, and 7 920 for E14.5). (B) Major fibroblast groups (details in Figure 2D-G and **Figure S2A**) and velocity trends (details in **Figure S2B**) overlayed on UMAP visualization of all fibroblasts (n = 25 944). (C) Dot plot depicting expression of unifying markers for major fibroblast groups. Highlighted are the clusters mostly present at E12.5. (D,F) Subclustering of major fibroblast groups (upper panel). Expression pattern of characteristic marker genes projected onto UMAP (lower panels). Maximum number of mRNA copies detected per cell is presented in brackets to provide an idea of the absolute abundance of the marker gene. (E,G) mRNA and protein stainings (RNA FISH in italics; protein IF in capital letters) revealing the anatomical location of fibroblast subpopulations. Dashed lines with arrows highlight the region with highest expression. Microscope images originate from larger tile scans (n = 3 mice). Scale bars, 100μm. (H) Expression pattern of characteristic marker genes projected onto UMAP revealing the subdivision of FIB Origin. (I) Absorption probabilities of the FIB Origin subpopulations towards the differentiation endpoints. Absorption probabilities projected onto UMAP (left panels) and absorption probabilities for each of the FIB Origin subpopulations (right panel). (J) mRNA staining revealing the spacial arrangement of FIB Origin subpopulations. Dashed lines with arrows highlight the region with highest expression. Microscope images originate from larger tile scans (n = 3 mice). Scale bars, 100μm. (K) Schemes summarizing major fibroblast groups and their location at E12.5.

**Figure 3.**
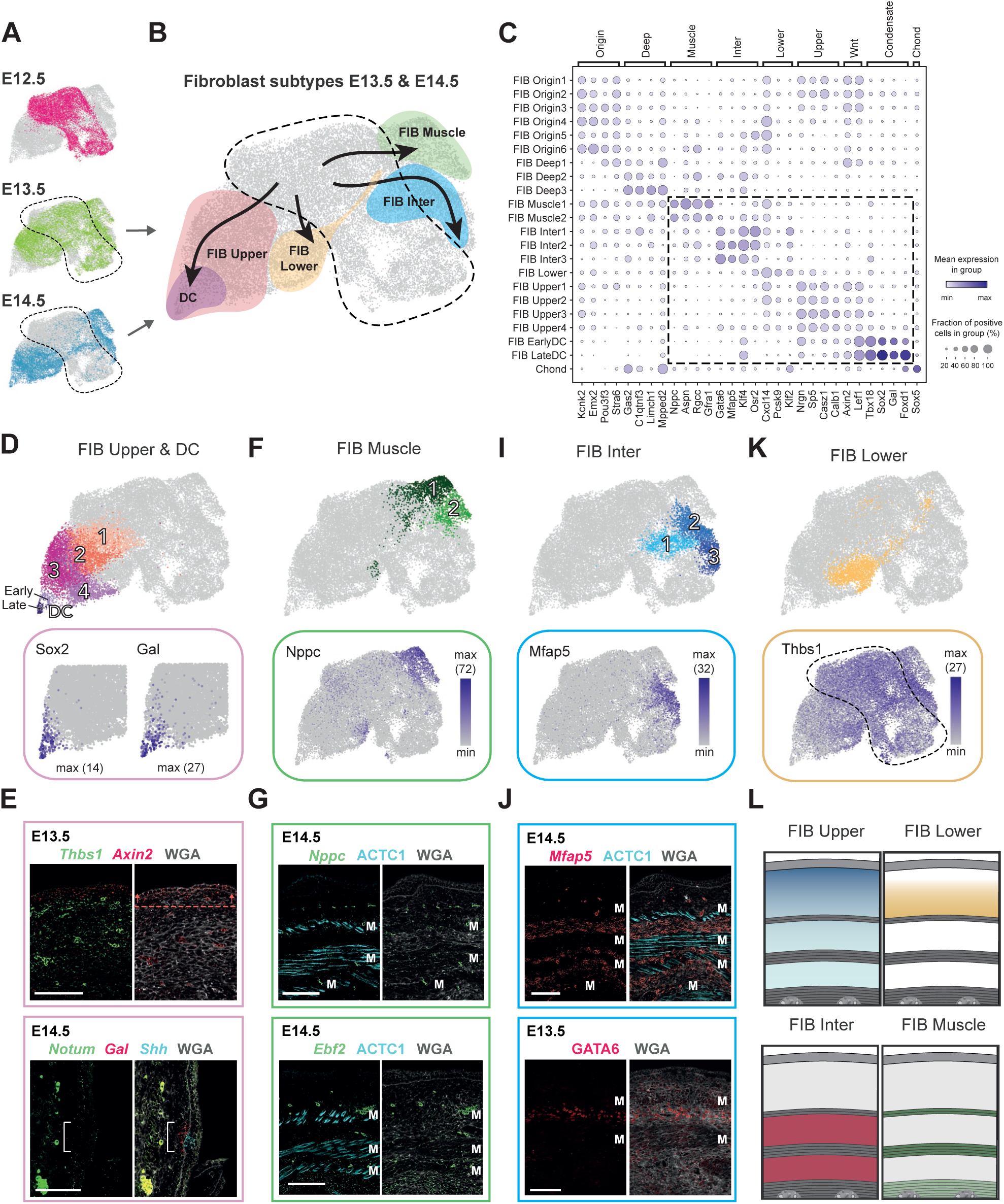
Deconstruction of fibroblast heterogeneity at E13.5 and E14.5 (expression and location) (A) UMAP visualization of all fibroblasts. Highlighted in color are the cells from the different embryonic ages (n = 10 008 cells for E12.5, 8 016 for E13.5, and 7 920 for E14.5). (B) Major fibroblast groups (details in Figure 3D-L and **Figure S2A**) and velocity trends (details in **Figure S2B**) overlayed on UMAP visualization of all fibroblasts (n = 25 944). (C) Dot plot depicting expression of unifying markers for major fibroblast groups. Highlighted are the clusters mostly present at E13.5 and E14.5. (D,F,I,K) Subclustering of major fibroblast groups (upper panel). Expression pattern of characteristic marker genes projected onto UMAP (lower panels). (E,G,J) mRNA and protein stainings (RNA FISH in italics, protein IF in capital letters) revealing the anatomical location of fibroblast subpopulations. Dashed line with arrows highlights the region with highest expression. Bracket highlights the region of reduced *Notum* expression. M marks developing muscle layers (for orientation in the tissue). Microscope images originate from larger tile scans (n = 3 mice). Scale bars, 100μm. (L) Schemes summarizing major fibroblast groups and their location at E14.5. E13.5 looks similar to E14.5 (with one muscle layer less).

### *FIB Origin* and *FIB Deep* – Notable fibroblast heterogeneity exists already at E12.5

At E12.5, dermis contains the two major fibroblast subsets *FIB Origin* and *FIB Deep* fibroblasts, characterized by expression of unique gene sets (**Figure 2A-C** and **S2C**) and different tissue locations (summarized in **Figure 2K**). The *FIB Origin* cells constitute a Wnt-pathway-activated *Lef1*^+^*Axin2*^+^ fibroblast subset that maps to the upper dermis (**Figure 2D,E**), and expresses remarkably few receptors and ligands (**Figure S2E**). According to RNAvelocity analysis, which can predict differentiation paths based on the expression of unspliced and spliced mRNA (La Manno et al., 2018), *FIB Origin* fibroblasts may serve as the source for almost all other fibroblast clusters emerging at E13.5 and E14.5 (**Figure S2B**). In addition to *FIB Origin* cells, there is a second *Lef1*^+^*Axin2*^+^ cell group at E12.5 *(FIB Deep1),* which is characterized by co-expression of *Pdgfc* (**Figure 2F**) and *Hoxb8* (**Figure S2F**) and maps to the lower half of the sub-epidermal space (**Figure 2G**). *FIB Origin* and *FIB Deep1* also share the expression of the myofibroblast markers *Acta2* (also known as aSMA) and *Tagln* (also known as SM22a) (**Figure S2G**). *FIB Deep2* and *FIB Deep3* constitute *Ebf2*^+^ *Postn*^+^ cell populations (**Figure 2F** and **S2H**), with *FIB Deep3* additionally expressing *Limch1* (**Figure S2F**). FIB *Deep2/3* were mapped within the deep back muscle at E12.5 (**Figure 2G** and **S2H**).

As the sub-epidermal/dermal tissue at E12.5 is still one compartment, which is not yet separated by any muscle layers (see **Figure 1M-O**), we considered that not all sampled fibroblasts (and/or their respective lineages) will be part of the skin-associated tissue compartment (i.e. fibroblasts above PCM, within PCM, and in fascia). Based on histological landmarks and tissue placement of *FIB Origin*/*Deep* subpopulations, *FIB Origin* cells were the most likely source for skin-associated fibroblasts. Thus, we probed whether *FIB Origin* cells are at E12.5 transcriptionally still uniform or show already heterogeneity that may point towards future fibroblast lineages. Unbiased clustering assigned *FIB Origin* cells into 6 subgroups (**Figure 2D**), which arranged in dimensionality-reduced space (UMAP) into “left” (*FIB Origin1/2*), “middle” (*FIB Origin3/4*) and “right” (*FIB Origin5/6*) subpopulations. The *FIB Origin1/2* cells are enriched for Wnt-pathway components such as *Lef1*, *Tcf7*, *Dkk1*, *Sp5* and the adhesion molecule *Cadm1* (**Figure 2C,D,H**), while the *FIB Origin5/6* cells are marked by e.g. *Epha4*, *Creb5*, and *Thbs2* (**Figure 2H**). *FIB Origin3/4* cells are in the UMAP placed between *FIB Origin1/2 and 5/6* cells and express genes of both. Co-staining of markers characteristic for *FIB Origin1/2* (*Dkk1-enriched*) or *FIB Origin5/6* (*Creb5-enriched*) fibroblast subsets revealed a clearly distinct spatial placement, with *FIB Origin1/2* mapping closer to epidermis than *FIB Origin5/6* (**Figure 2J**). Notably, when overlaying the marker gene expression for *FIB Origin1/2* and *Origin5/6* subsets on the fibroblast UMAP, the subsets seem to extend into *FIB Upper/DC* and *FIB Inter/Muscle*, respectively (**Figure 2H;** see **Figure 3B**), suggesting that the subdivision of *FIB Origin* cells may already reflect early commitment towards future fibroblast fates (presented in Figure 3). Indeed, fate simulation of the *FIB Origin* subpopulations confirmed this observation at a global gene expression level (**Figure 2I** and **S2I** and **Methods**). In summary, we found that skin-associated fibroblast heterogeneity already exists at E12.5 (*FIB Origin* subsets), likely reflecting their respective early fate specialization into functionally different fibroblast subsets. This represents the earliest reported fibroblast heterogeneity – transcriptionally and spatially – during the development of mouse skin.

### *CHOND* – Embryonic chondrocytes transcriptionally map with skin fibroblasts

Among the 22 fibroblast subsets we identified a small cluster (*CHOND*) that located next to the *FIB Deep* populations in the UMAP (**Figure S2A,J**), and expressed unique marker genes compared to all other fibroblasts (e.g. *Sox5, Sox6, Sox9, Col2a1, Col9a3, Acan*, *Matn4* and *Mia*) (**Figure S2K**). In-depth literature search revealed that this population constitutes chondrocytes (or chondrocyte precursors). Chondrocytes share their developmental origin (paraxial mesoderm-derived somites) with skeletal muscle and the dermis (Barresi and Gilbert, 2020). Developing cartilage cells express the transcription factors *Sox5* and *Sox6* that subsequently activate the cartilage-promoting factor *Sox9*, which then results in expression of chondrocyte-specific genes such as *Col2a1*, *Col9a3*, *Acan*, and *Matn4* (Yamashita et al., 2012; Zuo et al., 2018). Also *Pdgfra*, one of the prominent features shared between the chondrocyte and the other fibroblast subsets (**Figure S2K**), was recently shown to be critical for chondrocyte progenitor formation (Bartoletti et al., 2020). mRNA staining for the chondrocyte differentiation marker *Mia* (Bosserhoff and Buettner, 2003) confirmed our cell-type assignment as *Mia* showed strong expression in the developing vertebrae (**Figure S2L-M**). The realization that chondrocytes may cluster with fibroblasts can benefit future studies looking at fibroblast-like cell populations.

### *FIB Upper* and *FIB DC* – Acute loss of Wnt inhibitors marks dermal condensate formation

Starting from E13.5, as expected for the time just prior to hair follicle induction, we identified a fibroblast subset (*FIB Upper*) that shows high Wnt-pathway activity (e.g. *Lef1* and *Axin2*) and can mature into the hair follicle-inductive dermal condensate (*FIB DC*) (**Figure 2D** and **3A-D**). This is in line with the accepted view that Wnt-signaling activated fibroblasts are a prerequisite for embryonic hair follicle development (Andl et al., 2002; Chen et al., 2012; DasGupta and Fuchs, 1999; Enshell-Seijffers et al., 2010; Fu and Hsu, 2013; Gupta et al., 2019; Mok et al., 2019; Tsai et al., 2014; Zhang et al., 2009). *Axin2* mRNA staining revealed that these Wnt-signaling activated fibroblasts become confined to only a few layers in the uppermost dermis at E13.5 (compare **Figure 2E** with **3E**), a pattern that has been noted before (Chen et al., 2012). In line with previous reports (Biggs et al., 2018; Gupta et al., 2019; Mok et al., 2019), we also detect that cells exit the cell cycle just prior to DC commitment (upregulation of G0/G1 genes in *FIB DC*, e.g. *Cdkn1a* and *Btg1;* **Figure S2D** and **S3A**) and start expressing *Sox2* when the morphologically recognizable DC is forming (E14.5) (**Figure 3D** and **S2C**).

Additionally, our data revealed a most striking and abrupt gene-expression change at both *initial* and *final* DC-lineage commitment, each signified by a sharp downregulation of Wnt-pathway inhibitors. At the *FIB Origin* to *FIB Upper* border (E12.5 to E13.5; initial commitment) we noted acute and permanent downregulation of *Dkk2* (**Figure S3A**). Indeed, *Dkk2* has been found to be absent in hair-bearing skin while being expressed in non-hairy skin (Song et al., 2018). Together with our data this suggests that the absence of *Dkk2* may be a key determinator for fibroblasts becoming competent to enter a DC fate.

As *FIB Upper* cells become more Wnt-pathway activated, they also upregulate *Dkk1* (**Figure 2D,H**). This parallel upregulation continues until DC-fated cells acquire *Sox2* (final commitment), when *Dkk1* expression drops acutely (Gupta et al., 2019; Mok et al., 2019). Strikingly, in our data we observe the same switch-like pattern with sharp downregulation at the border between *FIB Upper3/4* and *FIB EarlyDC* also for *Notum* and *Cav1* (**Figure S3A**), both acting as Wnt-signaling inhibitors (Galbiati et al., 2000; Pentinmikko et al., 2019). The loss of *Notum* expression in mature DC cells was confirmed by co-staining of *Notum* and the DC-marker *Gal* in E14.5 skin (**Figure 3E**). The fact that several prominent Wnt-signaling inhibitors are first up-regulated in *FIB Upper* and then abruptly down-regulated in *EarlyDC* cells suggests that this is a functional feature of DC formation and/or DC commitment, which remains an exciting route to be explored.

### *FIB Lower* – Dermal fibroblasts without unique marker gene expression

At E13.5 another fibroblast subset (*FIB Lower*) emerges, which lacks unique marker gene expression. Thus, we placed the *FIB Lower* population via exclusion criteria within the tissue. At E13.5, *Axin2* expression becomes confined to the uppermost dermal fibroblasts (*FIB Upper*). *Thbs1* expression is concomitantly lost in those cells, but it is retained in the lower dermis, as well as in the subcutaneous interstitial layer below the PCM (**Figure 2E** and **3E,L** and **S3J**). The interstitium, illustrated in **Figure 1O**, is a fibroblast-and-fluid-filled space that starts below the PCM and reaches until the spine (interrupted only by the superficial back muscle layer) (Benias et al., 2018; Merrick et al., 2019). As subcutaneous interstitium (*Thbs1*^+^/*Mfap5*^+^; see below) can be reliably mapped back using *Mfap5* mRNA staining (**Figure 3J**), and *FIB Lower* cells do not express *Mfap5* (**Figure 3I**)*, FIB Lower* cells were assigned to the lower dermis.

### *FIB Muscle* – Perimuscular *Nppc*^+^ fibroblasts possess the ability to support the developing muscle

Also at E13.5, a group of *Nppc^+^* fibroblasts (*FIB Muscle*) was first observed (**Figure 3B** and **S2C**). This cell population is characterized by expression of *Nppc, Rgcc* and *Gfra1 (***Figure 3C,F**) and is exclusively located within the developing muscle layers (**Figure 3G**, upper image). This spatial confinement suggests a potential function of *FIB muscle* fibroblasts in supporting the developing muscle, which to date is unexplored. From work in adult murine skin it is known that fibroblasts are required for maintaining the balance between quiescence and activation of muscle stem cells (Murphy et al., 2011), and that they are the primary providers of extracellular matrix (ECM) to skeletal muscles (Gatchalian et al., 1989; Gillies and Lieber, 2011; Kühl et al., 1984; Sanderson et al., 1986; Sasse et al., 1981). Muscle ECM mediates force transmission from the fibers to the tendons and is dominated by the collagen isoforms I, III, IV, and VI (Cescon et al., 2015; Kjær, 2004; Light and Champion, 1984; Pace et al., 2008; Purslow, 2002; Tidball, 1991). Indeed, *FIB Muscle* cells express all these collagens at elevated or high levels (**Figure S3D**).

Unbiased clustering further separated *FIB Muscle* cells into two subgroups, with potentially distinct yet unknown functions. The two subgroups are characterized by expression of *Aspn* and *Wnt4* (*FIB Muscle1)* and *Ebf2* and *Igfbp3 (FIB Muscle2),* respectively (**Figure 3F** and **S3C**). As Wnt4 has been reported to maintain satellite cell quiescence (Eliazer et al., 2019), while Igfbp3 is known to support myoblast differentiation (Foulstone et al., 2003), it is conceivable that the two *FIB Muscle* subpopulations are involved in balancing activation and quiescence of satellite cells.

### *FIB Inter* – Fibroblasts constituting fascia fibroblasts and serving as a cellular source for adipose stem cells

From E13.5, a distinct group of *Mfap5+/Gata6+* fibroblasts (*FIB Inter*) can be detected (**Figure 3B,C,I**), which via mRNA staining for *Mfap5* – an ECM component (Kolehmainen et al., 2008) – could be mapped to the interstitial layers (**Figure 3J**).

When analyzing *FIB Inter* cells in more detail, we noticed that a subset, predominantly *FIB Inter1/2,* expressed genes that are characteristic for the fascia (**Figure S3F,G**), a connective tissue sheet that is located just below the PCM (Rinkevich et al., 2015). These genes include the most prominent fascia markers *Nov, Dpp4* and *Plac8* (Joost et al., 2020; Rinkevich et al., 2015), and additional fascia-associated genes such as *Mfap5, Wnt2, Creb5, Col14a1, and Tmeff2* (**Figure 3I** and **S3F,G**) (Joost et al., 2020).

In addition to fascia-associated fibroblast, *FIB Inter* also includes a subgroup of cells (*FIB Inter3*) that expresses the key adipogenic transcription factors *Pparg* and *Cebpa* (**Figure S3H** and **Table S1**) (Freytag et al., 1994; Tontonoz et al., 1994). Given that bundles of fascial fibres are often found mixed with fat to form preassure-tolertant fibro-adipose tissue associated with skin (e.g. soles, palms, or finger tips in humans) (Benjamin, 2009), we followed up on the intriguing possibility that *FIB Inter* cells might include adipogenic cells. We looked for early key regulators of adipogenesis such as *Junb*, *Fos*, *Atf3* and *Klf4* (Distel et al., 1987); all of which were expressed in *FIB Inter2* (**Figure S3H**). In contrast, *Cebpa* (Freytag et al., 1994) and *Pparg* (Tontonoz et al., 1994), which are transcription factors that drive later steps of adipogenesis, had their highest expression in *FIB Inter3* (**Figure S3H**), suggesting that adipogenesis progresses as cells mature from *FIB Inter2* to *FIB Inter3.* The terminal differentiation markers such as *Fabp4* and *Cd36* (Ferrero et al., 2020) are essentially absent (**Figure S3H**), which is in line with *Fabp4*^+^ dermal cells only appearing at E16 and the characteristic lipid droplets of mature adipocytes appearing at E18.5 (Wojciechowicz et al., 2013). In sum, our data suggests that a fibroblast subset as early as E13.5 may already be fated for adipose-lineage differentiation.

### Lineage tracing confirms *FIB Muscle* and *FIB Inter* populations

While the development of hair follicle associated fibroblasts, including lineage development and respective gene signatures, has been documented (e.g., Biggs et al., 2018; Ge et al., 2020; Gupta et al., 2019; Mok et al., 2019), not much is known about other functionally specialized fibroblasts such as muscle-associated or interstitial fibroblasts. Thus, we mined our data for the expression of genes that could be used for lineage tracing of *FIB Inter* and *FIB Muscle* cells. Our goal was not only to complement our expression and spatial data, but also to test whether *FIB Muscle* cells remain muscle restricted, and *FIB Inter* cells indeed contribute primarily to lower skin layers such as the fascia and adipose tissue as the scRNA-seq data suggested.

For tracing the *FIB Inter* cells, we identified *Gata6* as one of the most specific cell markers among all skin populations (at E13.5 almost exclusively expressed in *FIB Inter1-3;* **Figure 4A-B**). We used *Gata6-EGFP-CreERT2/R26-tdTomato* mice (hereafter called Gata6-Tom) and traced the fate from E13.5 to E15.5 (initial tracing), and from E13.5 to postnatal days P5 and P35 when hair follicles are in active growth (anagen), which is accompanied with an enlarged and mature adipose (DWAT) compartment (**Figure 4C**). At E15.5 adipocytes are not yet formed, however we found Gata6-Tom traced cells abundantly present in the fascia and subcutaneous interstitium (**Figure 4D**). Moreover, tracing at P5 and P35 revealed persistence of the *FIB Inter* lineage in the fascia from postnatal to early adulthood (**Figure 4E-F**). Unexpectedly however, we did not detect Gata6-Tom traced DWAT adipocytes; the presence of which we confirmed with PLP1 staining, a lipid-droplet associated protein that marks mature adipocytes (**Figure 4E-F**). Due to technical limitations subcutanous white adipose tissue (SWAT) was lost when harvesting postnatal skin. This limitation leaves open two possibilities; *FIB Inter* cells either do not represent adipocyte precursers or *FIB Inter* cells only contribute to SWAT formation and thus DWAT and SWAT originate from independent precursors (see **Discussion**).

**Figure 4.**
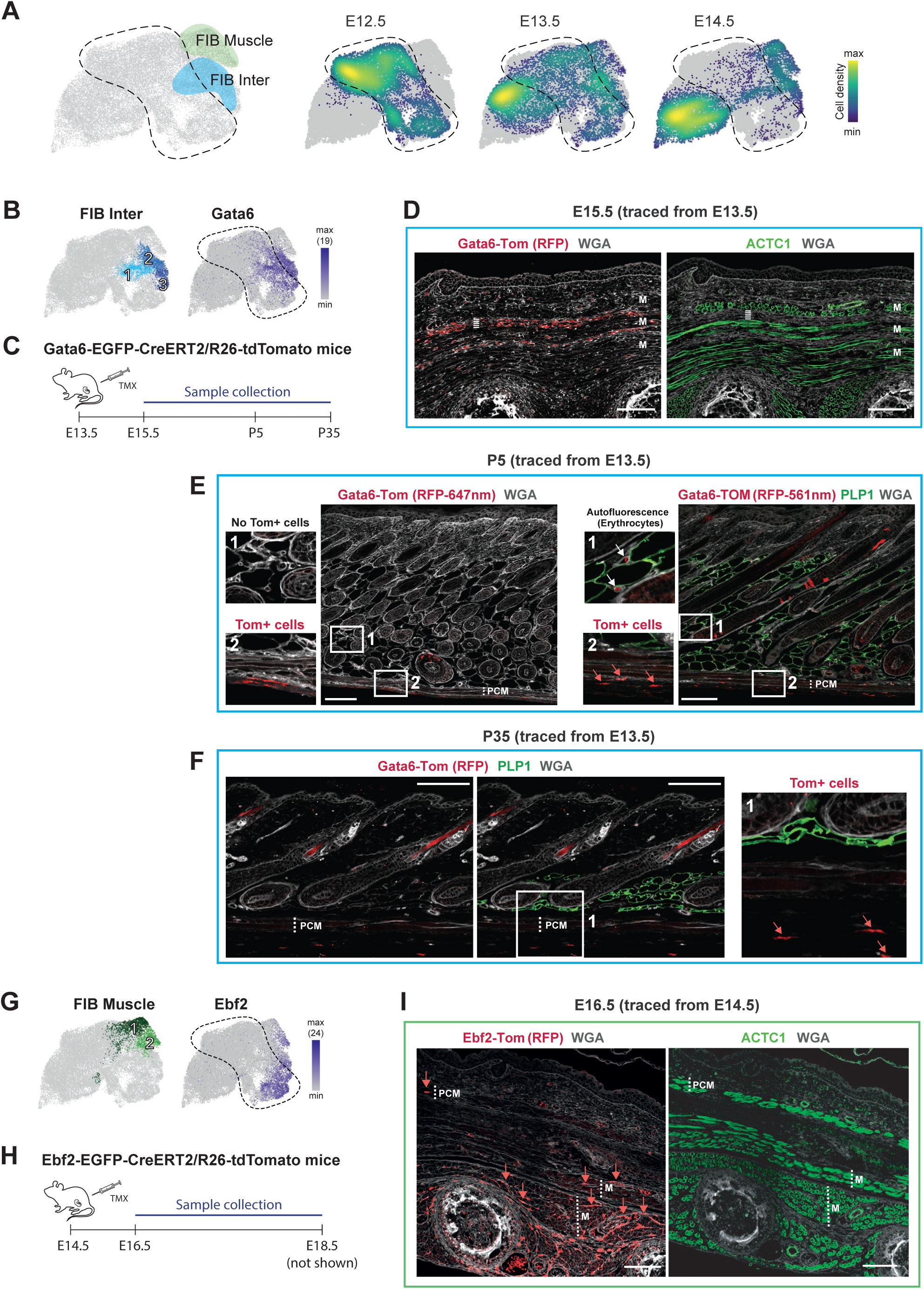
Tracing the fate of Ebf2+ and Gata6+ fibroblasts. (A) *FIB Muscle* and *FIB Inter* fibroblast groups (details in Figure 3D-K and **Figure S2A**) highlighted on UMAP of all fibroblasts (left panel). Density plot showing the distribution of fibroblasts from the different embryonic ages on the UMAP (right panel). (B) *FIB Inter* subpopulations projected onto UMAP (left panel). Expression pattern of *Gata6* (used for tracing) projected onto UMAP (right panel). (C) Experimental setup for lineage tracing of *Gata6*^+^ cells. (D) Initial 2-day-tracing pattern of *Gata6*^+^ cells stained with RFP antibody (left panel). ACTC1 protein staining of the developing muscle layers (right panel). Dashed line marks Fascia/SWAT layer (F/S) on the two consecutive sections. M marks developing muscle layers (for orientation in the tissue). Scale bars, 100μm. (E) Tracing pattern of *Gata6*^+^ cells at postnatal day 5 (P5) stained with RFP antibody, detected with 647-conjugated or 561-conjugated secondary antibody. PLP1 protein staining of lipid droplets. Note the strong erythrocytes autofluorescence at 561nm detection, which gives a false-positive image of DWAT being *Gata6*-Tom traced. Dotted lines indicate the PCM. Scale bars, 100μm. (F) Tracing pattern of *Gata6*^+^ cells at postnatal day 35 (P35) stained with RFP antibody. PLP1 protein staining of lipid droplets. Dotted lines indicate the PCM. Scale bars, 100μm. (G) *FIB Muscle* subpopulations projected onto UMAP (left panel). Expression pattern of *Ebf2* (used for tracing) projected onto UMAP (right panel). (H) Experimental setup for lineage tracing of *Ebf2*^+^ cells. (D) 2-day-tracing pattern of *Ebf2*^+^ cells stained with RFP antibody (left panel). ACTC1 protein staining of the developing muscle layers (right panel). Dotted lines indicate the PCM, underlying back muscle layers (M), as well as deep-tissue interstitial space on the two consecutive sections (for orientation in the tissue). Scale bars, 100μm.

To trace the *FIB Muscle* population, *Ebf2* was one of the most suitable markers (at E14.5 expressed in *FIB Muscle2* and *FIB Inter3;* **Figure 4A,G**). Thus, we utilized *Ebf2-EGFP-CreERT2/R26-tdTomato* mice (hereafter called Ebf2-Tom) and traced the fate of *Ebf2*^+^ cells from E14.5 to E16.5 and E18.5, respectively (**Figure 4H**). Both 2-and 4-day tracing gave rise to Ebf2-Tom cells within the deep back muscles and more rarely in the PCM, suggesting that the scRNA-seq-identified *FIB Muscle1/2* cell group indeed constitutes a muscle associated fibroblast subtype (**Figure 4I**). As expected from the scRNA-expression pattern (**Figure 4A,G**), Ebf2-Tom tracing also gave rise to some interstitial cells (**Figure 4I**).

### Cellular heterogeneity of vascular, immune, muscle and neural crest-derived cells in developing dermis

To determine potential supporting functions and interactions of fibroblasts with other major cell types in embryonic skin, we performed a detailed analysis of cellular heterogeneity among vessel-associated cells, immune cells, muscle cells and neural crest-derived cells (**Figure 5**) and further mined the data for potential interactions between the different cell populations showcasing a complex molecular interplay (**Table S3**). To identify receptor-ligand (R-L) pairs with likely involvement in vascular development, neuronal development, or immune cell recruitment, respectively, we manually scored each R-L-pair based on gene annotations and expression pattern (**Figure S5, Table S3** and **Methods**). Naturally, this approach did not uncover completely novel players, but it allowed us to rigorously subset for biologically relevant pairs, which overall broadened our analysis beyond the usual suspects and revealed expression patterns across all major cell types (**Figure 6**). Reassuringly, there was a significant overlap with previously identified signaling pairs in embryonic skin (Jin et al., 2021).

**Figure 5.**
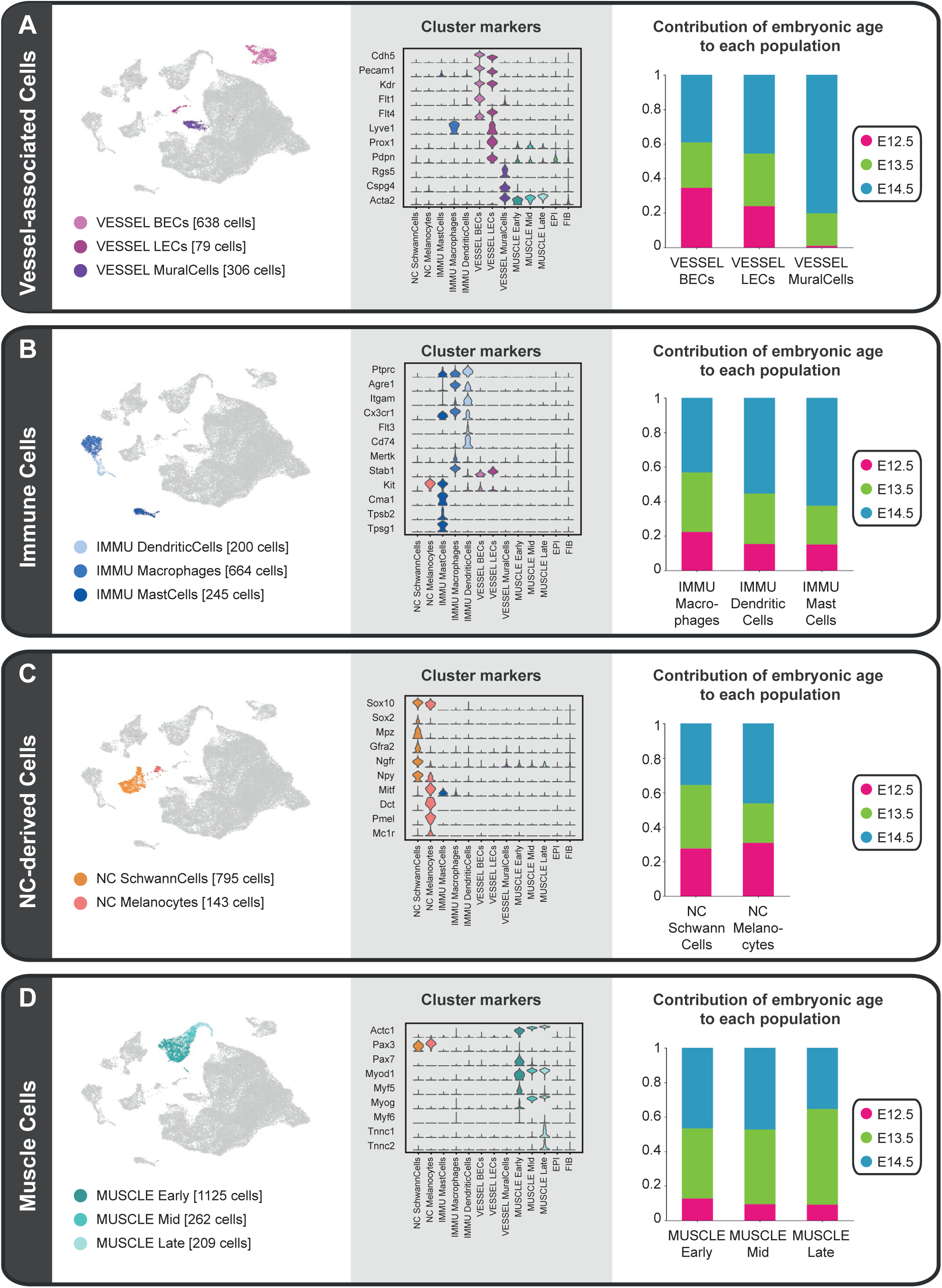
Cell types contributing to embryonic skin besides fibroblasts and keratinocytes. (A-D) Subpopulations of vessel-associated cells (A), immune cells (B), neural crest-derived cells (C), and muscle cells (D). Left panels: subpopulations projected onto UMAP from Figure 1B. Cell numbers per subpopulation are displayed in square brackets. Center panels: Violin Plot showing expression of marker genes that were essential to cell type assignment. Right panels: Bar plot visualizing the contribution of each embryonic time point to each subpopulation

**Figure 6.**
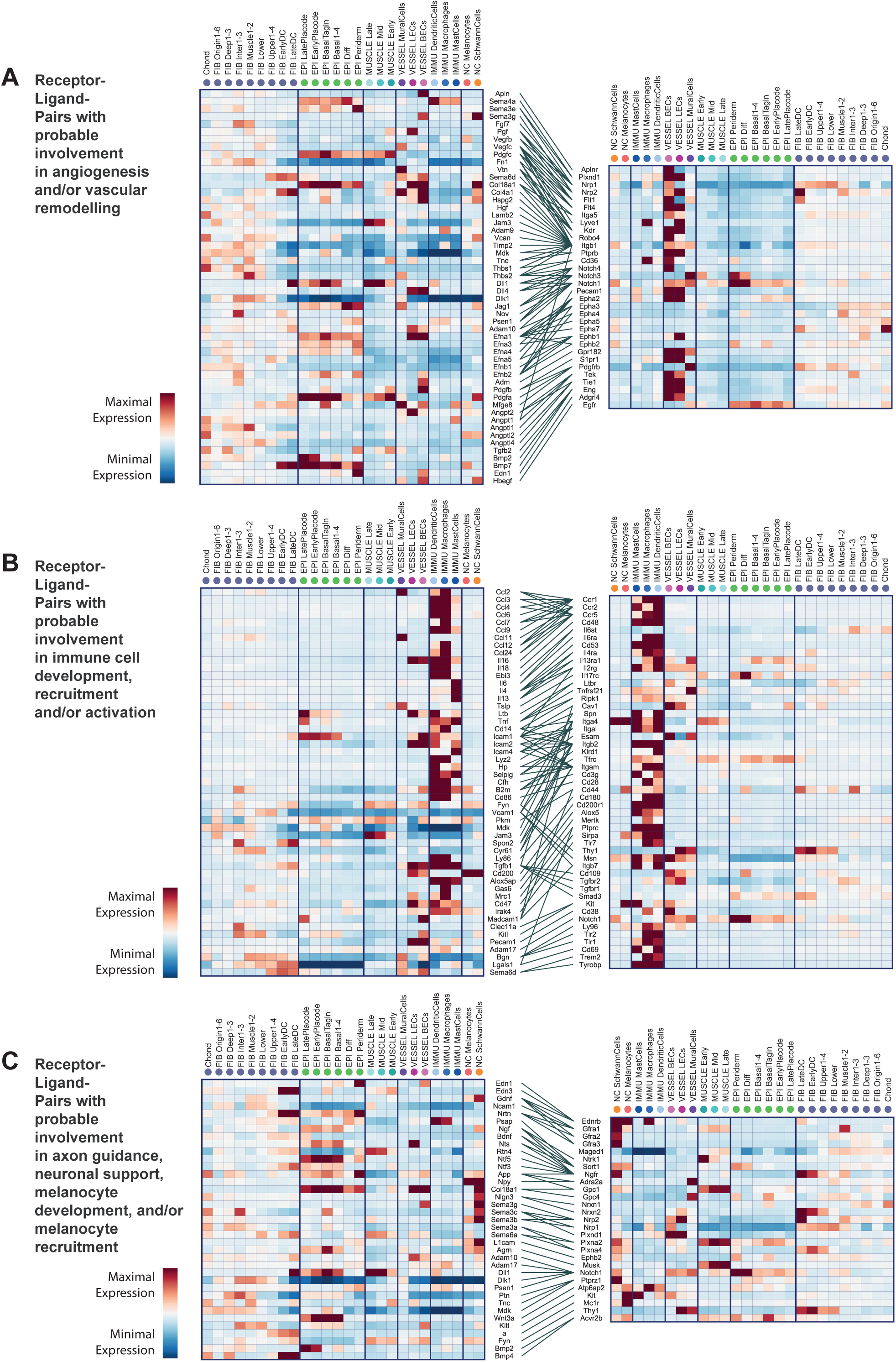
Interactions among all stromal and epithelial cell types. (A-C) Receptor-ligand pairs with probable involvement in angiogenesis and/or vascular remodelling (A), immune cell development, recruitment and/or activation (B), and axon guidance, neuronal support, melanocyte development, and/or melanocyte recrtuitment (C). Details on choice of included pairs are presented in **Figure S5**. Left and right panel: Heatmap showing the expression pattern of interaction partners among all first-level clusters as well as fibroblast and keratinocyte subclusters (partially grouped). Center panel: lines mark significant receptor-ligand pairs that were observed in our dataset. Left column contains mostly ligands, while right column contains mostly receptors. Note that *Icam1-Il2rg* and *Vcam1-Msn* (in A) and *Cd14-Itga2*, *Cd14-Itgb2*, *Cd14-Ripk1* (in B) do not strictly represent a receptor-ligand relationship, but rather two interacting receptors. Interaction partners were ordered manually to minimize crossing lines in the center panel. For details on presented interaction pairs also check **Table S3**.

### Vascular remodeling starts around E13, and sprouting vessels form without pericyte coverage in dorsal skin

In our dataset we detected vessel-associated cells including vascular and lymphatic endothelial cells (*VESSEL BECs* and *VESSEL LECs*, respectively), as well as vessel-surrounding mural cells (*VESSEL MuralCells*) (**Figure 1B-C** and **5A**).

To visualize the developing vasculature including blood and lymph vessels in the skin tissue (**Figure 1G-I**), we stained for *Pecam1* (better known as CD31) which is a pan-endothelial marker (Kriehuber et al., 2001). Reflecting that developing skin is in need of nutrients and oxygen, our *in situ* stainings revealed a dense cutaneous vascular network at each of the analyzed time points, which is in agreement with our sequencing data that identified *VESSEL BECs* and *VESSEL LECs* at all analyzed time points (**Figure 5A**). *VESSEL BECs* and *VESSEL LECs* can be well distinguished by differential expression of VEGF receptors, such as *Flt1* (also known as VEGFR1) only implicated in vascular angiogenesis, *Flt4* (VEGFR3) exclusively implicated in lymph angiogenesis, and *Kdr* (VEGFR2) implicated in both processes (Detmar, 2000) (**Figure 5A**).

*The VESSEL BECs* population encompasses cells that express *Efnb2*, *Ephb4*, *Apln*, and *Aplnr* (Apelin receptor), respectively (**Figure S4A**), suggesting that we captured both arterial and venous/capillary ECs. Jointly, these genes ensure proper alignment of arteries and veins for which an intricate balance of repulsion (via arterial *Efnb2* and venous *Ephb4)* and attraction (via arterial *Apln* and venous *Aplnr*) is required (Kidoya et al., 2015; Poliakov et al., 2004).

*VESSEL LECs* express a number of genes, e.g., *Lyve1, Prox1,* and *Pdpn* (**Figure 5A**), which are involved in the formation of lymphatic endothelium from venous endothelium (Alitalo et al., 2005; Ordóñez, 2012; Wigle and Oliver, 1999). That we observe *VESSEL LECs* already at E12.5 matches earlier work which reported murine dorsal skin to be invaded by a lymphatic network around E12.5 (Betterman and Harvey, 2018).

At E12.5, 13.5 and 14.5, vessels in skin are mainly formed by angiogenesis (Ribatti et al., 2000), which is well reflected in our data. At all analyzed time points, a significant fraction of our *VESSEL BECs* and *VESSEL LECs* expresses *Mmp2* and *Dll4* (**Figure S4A**), which signify ongoing angiogenesis. Sprouting vessels are led by so-called tip cells that express matrix metalloproteinases such as *Mmp2* to degrade the vascular basement membrane, which usually prevents endothelial cell migration, and the Notch ligand *Dll4* to prevent neighboring *Notch1^+^/ Notch4^+^* cells from responding to the key angiogenic factor *Vegfa* (Betsholtz, 2018; Mignatti and Rifkin, 1996; Ruhrberg et al., 2002).

Angiogenesis occurs in response to a large variety of angiogenic factors, such as VEGF family members, PDGF family members, BMPs, *Tgfb1*, *Pgf*, ECM components (e.g., *Pcolce*, *Col1a1*, *Sparc*), and molecules better known from axon guidance (Semaphorins, Netrins, Neuropillins, Slit proteins) (Cao et al., 2002; David et al., 2009; Detmar, 2000; Goumans and Mummery, 2000; Luttun et al., 2002; Newman et al., 2011; Shima and Mailhos, 2000). Our data shows that these factors are provided by a variety of different cell types (**Figure S4A**) and that fibroblasts expressed the broadest spectrum of typical angiogenic growth factors (**Figure 6A**). Although we could not locate the cellular source of *Vegfa* (we only detected *Vegfa* in very few cells, mostly fibroblasts), it is interesting to note that the fibroblasts seem to collectively provide angiogenic factors as the expression pattern did not identify particular fibroblast subpopulations to be predominantly in charge (**Figure 6A**). Notably, also the epidermis expresses high levels of angiogenic factors such as *Pdgfa, Bmp2*, or *Bmp7* (**Figure 6A**), which fits the earlier notion that avascular epidermis might provide angiogenic stimuli to enhance dermal blood supply and match its metabolic needs (Malhotra et al., 1989).

For full vascular functionality and integrity, angiogenesis has to be followed by vascular remodeling, i.e. the recruitment of mural cells (Levéen et al., 1994; Soriano, 1994). Mural cells encompass vascular Smooth Muscle Cells (vSMCs) and pericytes, which we presented as one *Rgs5^+^/Cpsg4^+^/Acta2^+^* population as their transcriptomes are too similar for unequivocal separation (Gerhardt and Betsholtz, 2003; Muhl et al., 2020). Mural cells are recruited by endothelial cells that secrete *Pdgfb* and *Hbegf* (in our data expressed by *VESSEL BECs*), which can be sensed by *Pdgfrb*^+^ *Egfr*^+^ mural cells (**Figure S4A**) (Stratman et al., 2010).

Our data suggests that in dorsal skin, vascular remodeling does not happen before E13.0. At E12.5 we only classified 3 cells from the entire dataset as mural cells (**Figure 5A**), and staining for *Rgs5* mRNA, a specific marker for mural cells, did not reveal *Rgs5* positive cells (**Figure 1G**). The number of mural cells clearly increased by E13.5 (**Figure 1H** and **5A**) and at E14.5, staining revealed small vessels with sparse lining of *Rgs5*^+^ cells (likely representing pericytes) and a few larger vessels with continuous lining of *Rgs5*^+^ cells (likely representing vSMCs) (**Figure 1H**; smaller vessel in upper detail and larger vessel in lower detail). The absence of pericytes at E12.5 is significant because of a longstanding controversy whether sprouting vessels initially form without pericyte coverage (Benjamin et al., 1998), or if pericytes are present from the beginning and actively assist vessel sprouting (Amselgruber et al., 1999; Reynolds et al., 2000). Our data clearly supports the first model for embryonic dorsal skin.

### Immature skin is already primed with mast cells, dermal dendritic cells and immature macrophages

Our data revealed that, between E12.5 and E14.5, dorsal skin is exclusively populated by myeloid cells, i.e. dermal dendritic cells (*IMMU DendriticCells*), macrophages (*IMMU Macrophages*) and mast cells (*IMMU MastCells*) (**Figure 1B,C** and **5B**). Each of these populations express *Ptprc* (better known as CD45), they are present at all analyzed embryonic time points (**Figure 5B**) and exclusively locate to the dermis (**Figure 1J**).

While dendritic cells and macrophages share the expression of a number of markers such as *Adgre1* (better known as F4/80), *Itgam*, and *Cx3cr1* (Haniffa et al., 2012; Henri et al., 2010), they clearly differ in their development as well as function, which is reflected in the differential expression profiles (**Figure 5B** and **S4B**). Dendritic cell development is critically linked to *Flt3* which is reflected in high, dendritic cell-specific *Flt3* expression (**Figure 5B**) (Merad and Manz, 2009). Functionally, they are specialized on inducing the adaptive immune system and thus are well equipped to capture, process and present antigens via MHC-II complexes (MHC-II related genes include e.g. *H2-Aa, H2-Ab1, H2-Eb1, Cd74* and the transcriptional master regulator *Ciita*) (**Figure 5B** and **S4B**) (Nussenzweig et al., 1980; Pierre et al., 1997). Furthermore, dermal dendritic cells express *Ccr2* and *Ccr7* which are essential for their migration to the skin-draining lymph nodes where they meet and activate T cells (**Figure S4B**) (Förster et al., 1999; Steinman et al., 1978). When it comes to dendritic cell subsets, our dendritic cell population contains mostly classical dendritic cells marked by *Itgam* expression (better known as CD11b) and a minority of classical dendritic cells marked by *Itgae* expression (better known as CD103) (**Figure 5B** and **S4B**) (Henri et al., 2010; Mildner and Jung, 2014; Tamoutounour et al., 2013). To our knowledge, it has not been reported when dendritic cells start seeding the mouse dermis; our data shows that they are already present at E12.5 (**Figure 5B**).

Dermal macrophages are tissue-resident cells that are specialized to scavenge damaged cells or invading bacteria (Hashimoto et al., 2011). *Mertk* and *Stab1*, two crucial receptors for the scavenging function (Gautier et al., 2012), are specifically upregulated in our macrophage population (**Figure 5B**). Skin macrophages acquire MHC-II expression during their maturation process, but the majority of macrophages detected in our dataset still lack MHC-II molecule expression and thus seem to mostly represent immature macrophages (**Figure S4B**). We detect macrophages at all analyzed time points, and indeed macrophage-like cells have been described to seed the dermis as early as E10.5 (Hoeffel et al., 2012, 2015). There is a possibility that our macrophage population also contains precursors of Langerhans Cells as those can derive from yolk sac-derived macrophages and share a number of molecular features with them (*Adgre1*^+^*, Ptprc*^+^*, Itgam*^+^*, Cx3cr1*^+^*, Flt3*^-^) (**Figure 5B** and **S4B**) (Chorro et al., 2009; Hoeffel et al., 2012; Ginhoux and Merad, 2010).

The third immune population in this dataset constituted mast cells (MCs). These were characterized through e.g. expression of *Kit* and by the presence of mediators such as serine proteases (e.g. *Cma1, Tpsb2, Tpsg1*), which are typical for Mast cell secretory granules (**Figure 5B**) (Dwyer et al., 2016). It was shown recently that the first-wave yolk sac-derived MCs are replaced by definitive MCs during late embryonic development (Gentek et al., 2018). Our MCs clearly display the signature of yolk sac-derived MCs (*Grm6*^+^*, Cma1*^+^*, Prss34*^+^*, Smpx*^+^), while lacking the definitive, adult MC signature (*Adrb2*^-^*, Il1rap*^-^*, C2*^-^*, Lyz1*^-^) (**Figure 5B** and **S4B**). That we find MCs already at E12.5 (**Figure 5B**) contrasts earlier publications reporting sparse dermal mast cells only at around E14.5/E15 (Gentek et al., 2018; Hayashi et al., 1985).

Leukocyte recruitment to peripheral tissues is directed by chemokines. Our receptor-ligand analysis again provides a valuable overview of the expression pattern of the plethora of chemokines involved in leukocyte recruitment (**Table S3** and **Figure 6B**). We found most striking, that mostly skin-resident immune cells (and partly mural cells) express those chemokines suggesting their active involvement in skin homing of more immune cells.

### Melanocytes and Schwann cells in early embryonic skin

Our dataset contains two neural crest-derived populations: Schwann cells (*NC SchwannCells*) and melanocytes (*NC Melanocytes*), which are sampled at all three time points and marked e.g. by *Sox10* expression (**Figure 1B,C** and **5C**) (Kuhlbrodt et al., 1998).

Peripheral neurons have entered embryonic skin at our studied time points (Jenkins and Lumpkin, 2017; Martin, 1990), which is reflected by the presence of *Sox2*^+^*, Mpz*^+^*, Gfra2*^+^ Schwann cells (Peng et al., 2020) in our dataset (**Figure 5C**). Yet, we did not pick up neuronal-cell transcriptomes likely reflecting that the neuronal cell bodies are located either in the spinal cord (in the case of motor neurons), in paravertebral sympathetic ganglia (sympathetic neurons), or in dorsal root ganglia (sensory neurons) (Frank and Sanes, 1991). By visualizing the nerve-encasing Schwann cells in skin tissue with *Sox10* mRNA and PPARG protein (Yamagishi et al., 2008) staining, we observed thick sensory nerve trunks traversing the dermis towards the epidermis at E12.5 (**Figure 1K** and **S4C**), which seem to split up and spread out as the embryo continues to grow (**Figure 1L**). At E14.5 the majority of nerves are located directly underneath the epidermis (sensory neurons) and under the PCM (sensory and motor neurons) (**Figure 1L** and **S4C**).

Proper innervation of skin is facilitated by neuronal as well as non-neuronal cells, such as fibroblasts and keratinocytes. They together express numerous neurotrophins crucial for neuron growth and maintenance (e.g. *Ntf3, Ntf5*, *Bdnf* and *Ngf*), as well as a variety of genes that direct the growth cones of developing axons (e.g. Ephrins, Netrins, Slit proteins and Semaphorins) (**Figure 6C** and **S4D**) (Allen and Dawbarn, 2006; Ansel et al., 1996; Botchkarev et al., 2006; Di Marco et al., 1991; Roggenkamp et al., 2012; Roosterman et al., 2006). In turn, cutaneous nerves also release a variety of neuropeptides, such as *Npy* (Neuropeptide Y), to increase vascular permeability, support immune cell recruitment, and induce angiogenesis (**Figure 5C**) (Movafagh et al., 2006; Paus et al., 2006).

The melanocytes in our dataset were identified through a set of genes that are essential for melanocyte development; among them the master melanocyte transcription factor *Mitf*, other specification markers such as *Dct*, *Pmel,* and the key melanogenic enzyme tyrosinase (*Tyr)* (**Figure 5C**) (Chang et al., 2014; Ostrowski and Fisher, 2018). It has been described that melanoblasts migrate laterally through the dermis around E12.5, and enter the epidermis around E13.5 (Mayer, 1973; Yoshida et al., 1996). This is what we also observe in our tissue stainings, where the majority of *Sox10*^+^*/Pmel*^+^ melanoblasts has not yet entered the epidermis at E12.5 (**Figure 1K**), while at E13.5 (data not shown) some melanoblasts already sit within the epidermis (**Figure 1M-O**). At E14.5, melanoblasts are spread throughout the whole epidermis (**Figure 1L**). It is only during early postnatal life that IFE-resident melanoblasts are lost and hair follicle-homed melanoblasts persist (Hirobe, 1984).

Notably and well-fitting with recruitment of melanoblasts to the epidermis, our epidermal cell populations as well as fibroblasts located closest to the epidermis indeed express genes like *Edn1*, *Edn3*, and *Kitl* (implied in melanocyte survival and migration (Chang et al., 2014)), as well as *a* (also known as *Agouti*; regulating melanogenesis via the melanocortin receptor 1 (*MC1r*) (**Figure S4D** and **6C**) (Suzuki et al., 1997)).

### Panniculus carnosus muscle transcriptionally resembles other back muscles

We identified muscle cells at all three sampled time points (**Figure 1A-C** and **5D**). Staining for ACTC1 protein (Actin alpha cardiac muscle 1), the predominant actin isoform in early muscle development (Hayward and Schwartz, 1986), reveals the different muscle layers traversing the embryonic back. While at E12.5 there are only the deep back muscles that connect the developing vertebrae, by E13.5 superficial back muscles have developed, and by E14.5 the PCM (the most superficial muscle layer just underneath the DWAT layer in adult murine skin) has appeared (**Figure 1D-F**). To our knowledge, the timing of PCM development has not been reported for dorsal mouse skin.

With our transcriptomic data, we captured the full process of early skeletal myogenesis (**Figure 5D**). The *MUSCLE Early* population contains *Pax7*^+^ skeletal muscle stem cells, so-called satellite cells (Relaix et al., 2006; Seale et al., 2000). The *MUSCLE Mid* and *MUSCLE Late* populations recapitulate the stepwise changes in expression of myogenic regulatory factors that govern muscle cell differentiation. *Myod1* and *Myf5* are early markers for satellite cells that have committed to differentiation, while *Myog* and *Myf6* drive proliferation arrest and terminal myoblast differentiation (**Figure 5D**) (Grzelkowska-Kowalczyk, 2016; Zammit et al., 2006). In the *MUSCLE Late* population we furthermore detected markers of mature skeletal muscle fiber subtypes, e.g. *Tnnc1* and *Tnnc2* for Type I and Type II fibers, respectively (**Figure 5D**) (Rubenstein et al., 2020; Schiaffino and Reggiani, 2011).

In our dataset, the majority of muscle cells were satellite cells (**Figure 5D**). This is in line with previous literature, reporting that the proportion of satellite cells drops with developmental progress. While in early postnatal life satellite cells still account for 30-35% of muscle cells, this proportion is reduced significantly to 1-4% in adult mice (Zammit, 2008). Moreover, as we capture each of the muscle subpopulations (*MUSCLE Early, Mid, Late*) at all sampled time points, we concluded that every subpopulation likely is comprised by a mixture of cells from the deep back muscles, the superficial back muscles, and the PCM. This is interesting as it suggests that there is no major difference between the different muscle layers at the transcriptomic level.

As with other skeletal muscles in the body, the PCM as well as the back muscles are innervated by motor neurons which connect to the muscle via neuromuscular junctions that are scattered along the myofibers (Holstege and Blok, 1989; Petruska et al., 2014; Theriault and Diamond, 1988). We indeed find evidence of those neuromuscular junctions in our data (**Table S3** and **Figure S4E**). In the *MUSCLE Late* population there is expression of *Musk* (Muscle Associated Receptor Tyrosine Kinase), that upon binding of *Agrn* (expressed in motor neurons and other neural crest-derived cells) contributes to the formation of neuromuscular junctions by inducing the clustering of acetylcholine receptors in the postsynaptic neuromuscular junction (Reist et al., 1992; Tan-Sindhunata et al., 2015). Subunits of those acetylcholine receptors (e.g. *Chrna1, Chrna4, Chrnd, Chrng*) are expressed in *MUSCLE Mid* and *MUSCLE Late* cells (**Figure S4E**).

So far, we have focused on the temporo-spatial maturation of the dermis, its developing cellular composition and molecular communication potential between cell types. Once the dermis is successively equipped with all necessary cell types and structures, it is ready to support the full epidermal maturation, including hair follicle induction and epidermal stratification.

### Basal epidermal heterogeneity starts already at E12.5

At E12.5, the epidermis consists of a morphologically uniform basal layer that is covered by the periderm. Surprisingly, the basal cells separate already at this developmental timepoint into two transcriptionally distinct populations, which we termed *EPI BasalTagln* and *EPI Basal1* (**Figure 7A-C** and **S6A,C**). *EPI Basal1* cells have no unique molecular signature within keratinocytes. Their signature consists of genes that are shared with either *EPI BasalTagln* from E12.5 skin (e.g., *Olfm1*, *Bfsp2*, *Acvr2a, Podxl*) or with *EPI Basal2–4* populations from E13.5 and E14.5 skin (e.g., *Lmo1*, *Dcn*, *Ifitm3*) or with all basal populations (e.g., *Krt5, Krt14, Krt15, Sostdc1, Vcan*) (**Figure S6B**). Based on gene expression and their position in dimensionality-reduced space it seems likely that *EPI Basal1* cells are the precursors for the general interfollicular epidermis (IFE) at E13.5 and E14.5 (**Figure 7A** and **S6A**).

**Figure 7.**
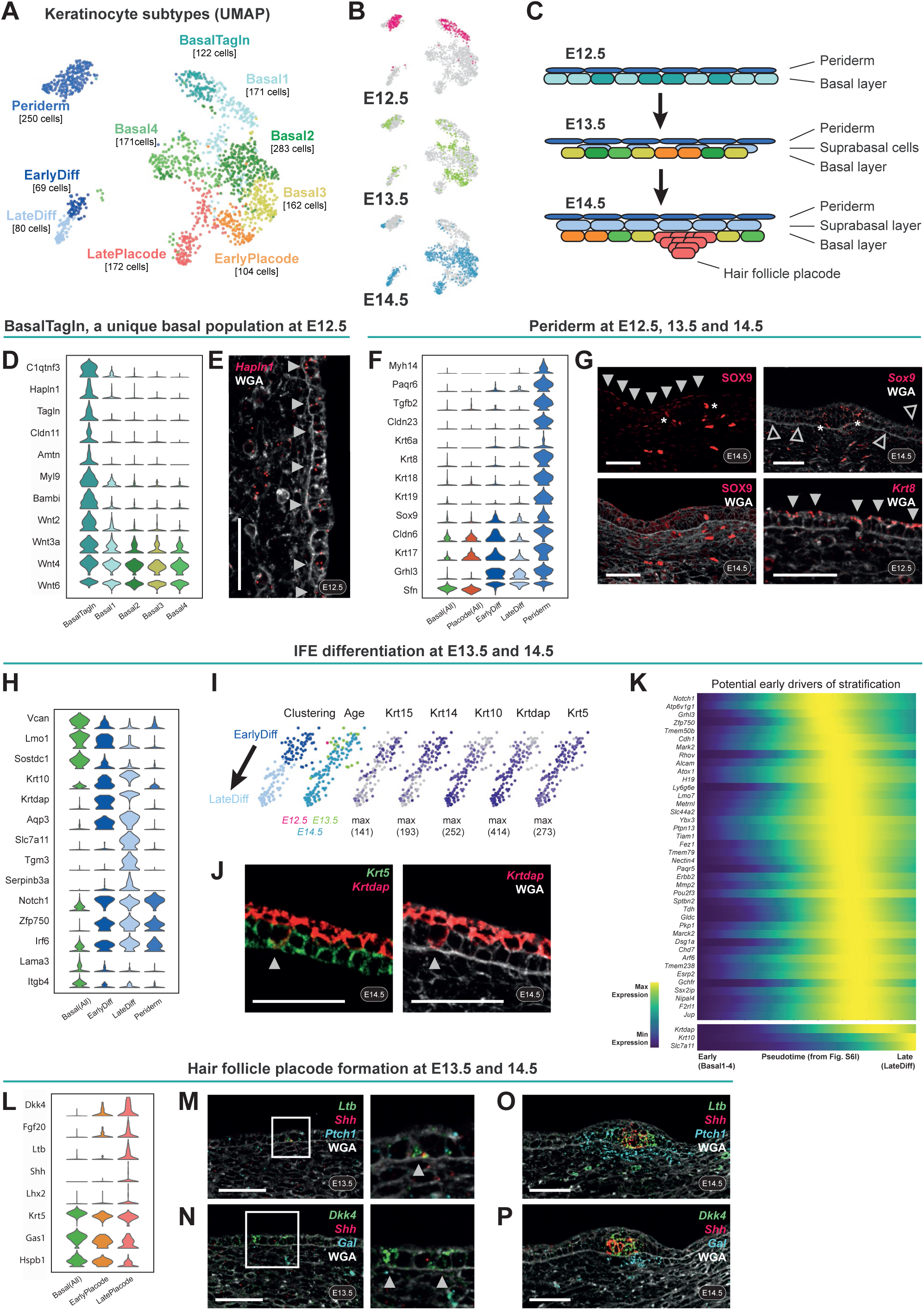
Epidermal development from a single basal layer towards a HF-inducing and stratified epithelium. (A) UMAP visualization of all keratinocytes, colored according to keratinocyte subclustering. Cell numbers per cluster are displayed in square brackets. (B) UMAP visualization of all keratinocytes. Highlighted in color are the cells from the different embryonic ages (n = 360 cells for E12.5, 347 for E13.5, and 877 for E14.5). (C) Scheme summarizing the development of the epidermis during the analyzed time window. Cells are colored according to the cluster colors in (A). (D) Violin Plot showing the cluster-specific expression of *EPI BasalTagln* marker genes. (E) *Hapln1* mRNA staining in dorsal E12.5 skin revealing that *Hapln1*^+^ (*EPI BasalTagln*) cells indeed map to the basal interfollicluar epidermis (arrowheads). Counterstained with WGA. Microscope image originates from larger tile scan (n = 3 mice). Scale bar, 50μm. (F) Violin Plot showing the expression of known and novel periderm marker genes. (G) SOX9 protein staining in dorsal E14.5 skin (left panels) highlights SOX9^+^ periderm cells (filled arrowheads) as well as SOX9^+^ cells in periphery of hair placode (asterisks). *Sox9* mRNA staining in dorsal E14.5 skin (upper right panel) shows accumulation of signal in periphery of hair placode (asterisks), but also sparse basal cells outside the placode that express *Sox9* mRNA (empty arrowheads). *Krt8* mRNA staining in dorsal E12.5 skin confirms *Krt8* as a specific periderm marker (filled arrowheads). Counterstained with WGA. Microscope images originate from larger tile scans (n = 3 mice). Scale bars, 50μm. (H) Violin Plot showing the gradual loss of basal markers and the concomitant gain of differentiation markers in differentiating keratinocytes, as well as a shared signature between differentiating keratinocytes and periderm cells. (I) Keratinocyte subclustering and embryonic age (left two panels) as well as expression pattern of basal and suprabasal marker genes (right five panels) projected onto part of the UMAP from (A) (only *EPI EarlyDiff* and *EPI LateDiff* cells). Cells are colored according to the cluster colors in (A) or embyronic age colors in (B), respectively. Maximum number of mRNA copies detected per cell is presented in brackets to provide an idea of the absolute abundance of the marker gene. (J) *Krt5* and *Krtdap* mRNA staining in dorsal E14.5 skin reveals a representative basal cell with a differentiation signature (arrowhead). Counterstained with WGA. Microscope image originates from larger tile scan (n = 3 mice). Scale bars, 50μm. (K) Heatmap of potential early drivers of stratification. A pseudotime was modeled based on *EPI Basal1-4*, *EPI EarlyDiff*, and *EPI LateDiff* cells from E14.5 (**Figure S6I**) and pseudotime-dependent genes were determined. Genes peaking earliest in pseudotime were chosen for display on heatmap. *Krtdap*, *Krt10*, and *Slc7a11* are plotted for comparison. (L) Violin Plot showing positive and negative marker genes of the hair placode. (M-N) *Ltb, Shh* and *Ptch1* mRNA staining (M) and *Dkk4*, *Shh*, and *Gal* mRNA staining (N) in dorsal E13.5 skin reveals early placode cells (arrowheads). Counterstained with WGA. Microscope images originate from larger tile scans (n = 3 mice). Scale bars, 50μm. (O-P) *Ltb, Shh* and *Ptch1* mRNA staining (O) and *Dkk4*, *Shh*, and *Gal* mRNA staining (P) in dorsal E14.5 skin reveals mature hair placodes (marked by *Ltb*, *Shh*, and *Dkk4*) as well as dermal condensates (marked by *Ptch1* and *Gal*). Counterstained with WGA. Microscope images originate from larger tile scans (n = 3 mice). Scale bars, 50μm.

In contrast, the *EPI BasalTagln* cells have a unique set of marker genes that is not shared with any other epidermal cell population. This unique gene set includes for example *C1qtnf3*, *Hapln1*, *Tagln*, *Cldn11*, *Amtn*, *Myl9*, *Bambi*, and many more (**Figure 7D-E** and **Table S1**). It was also remarkable to find that *EPI BasalTagln* cells express smooth muscle genes such as *Tagln* (also known as SM22a) and *Myl9* (**Figure 7D**), which is indeed unexpected for an epithelial cell population under physiological conditions. Compared to all other basal cell clusters, *EPI BasalTagln* cells express a high number of receptors and even more ligands, including unique expression of *Wnt2* and significant upregulation of *Wnt3a, Wnt4* and *Wnt6* (**Figure 7D, S6E,** and **Table S1**) – which were commonly believed to be expressed uniformly throughout embryonic epidermis (Andl et al., 2002; Reddy et al., 2001).

In contrast to significant heterogeneity in E12.5 basal IFE, basal IFE at E13.5 and E14.5 appears transcriptionally rather uniform. Basal IFE subpopulations *EPI Basal2-4* from E13.5 and E14.5 skin only show minor differences in expression (many of which are related to cell cycle) and we did not identify unique gene signatures (**Figure S6B** and **Table S1**). It furthermore stands out, that basal IFE at E13.5 and E14.5 is hardly expressing any specific receptors or ligands (**Figure S6E**). This is somewhat surprising as the dramatic epithelial changes including epidermal stratification and placode formation start at E13.5 and one would expect extensive inductive signaling to be ongoing at E13.5.

### The periderm matures and exhibits a surprisingly signaling-rich molecular signature

Periderm is a curious specialization of embryonic epidermis. It is a layer composed of squamous cells that cover the epidermis and its presence is crucial for preventing pathological epithelial adhesions within the embryo. Periderm cells start covering dorsal epidermis around E10 (they delaminate from the basement membrane and migrate up to cover basal keratinocytes) and are shed around E17/18 when the cornified layer forms (Hammond et al., 2019; M’Boneko and Merker, 1988; McGowan and Coulombe, 1998; Richardson et al., 2014). Although the existence of this layer has long been known, its molecular characterization has remained incomplete. In previous scRNA-seq studies of embryonic mouse skin, the periderm cells did not form their own cluster; instead they were interspersed among other keratinocyte populations (e.g., reanalysis of (Gupta et al., 2019); data not shown). In our dataset, we robustly identify the periderm cells as their own cluster (‘*EPI Periderm’*) (**Figure 7A** and **S6A**) likely due to the larger cell proportion relative to all epithelial cells at E12.5 (**Figure 7B** and **S6C**). Present at all three time points, we identified new (e.g. *Myh14, Paqr6, Tgfb2, Sox9*) as well as known periderm markers (e.g. *Cldn6/23, Krt6/8/17/18/19, Grhl3, Sfn*) (Dale et al., 1985; De La Garza et al., 2013; McGowan and Coulombe, 1998; Morita et al., 2002; Richardson et al., 2014), of which *Cldn6, Krt17, Grhl3*, *Sfn and Sox9* are not uniquely expressed in *EPI Periderm* cells but shared with selected other keratinocyte populations (**Figure 7F** and **Table S1**). Interestingly, *Sox9* and *Krt8,* which acclaimed fame in the context of hair follicle formation and as a marker of the early immature epidermis, respectively, show their highest mRNA expression in periderm cells (**Figure 7F-G**). Regarding *Sox9* expression itself, we additionally observed a similar basal-layer (placode periphery) and suprabasal (IFE associated) pattern as recently reported (**Figure 7G**) (Morita et al., 2021).

Cells within the periderm layer still undergo mitotic divisions similar to differentiated IFE cells (Damen et al., 2021; Lechler and Fuchs, 2005; Richardson et al., 2014), even though these cells have delaminated (**Figure S6D**). In addition, we found that the periderm undergoes molecular maturation characterized by increased expression of known IFE differentiation markers (e.g., *Krtdap*, *Lgals3*, *Dkkl1*), and multiple genes that have not been linked to epidermal differentiation, such as *Foxq1*, *Krt4*, *Lingo2*, *Mal*, *Pllp*, *Prss27*, and *Tchh* (**Figure S6F**). The function of these genes in periderm remains to be investigated, but it is conceivable that upon epidermal stratification the periderm starts to slowly prepare for its shedding. Most surprising however was that among all keratinocyte clusters *Periderm* cells express the highest number of receptors and ligands (**Figure S6E**), including those facilitating e.g., Ephrin signaling, Notch signaling, Igf signaling and Tgfb signaling (**Table S1**).

### Deconstructing early epidermal stratification

Differentiating keratinocytes separate into an early differentiating group (called *EPI EarlyDiff*) marked by co-expression of basal (e.g., *Krt15, Krt14*, *Vcan*, *Lmo1, Sostdc1*) as well as suprabasal genes (e.g., *Krt10*, *Krtdap*, *Aqp3*), and a more mature differentiating population (*EPI LateDiff*) which has gradually lost basal gene expression, further increased expression of suprabasal genes and acquired the expression of more mature differentiation markers (e.g., *Slc7a11, Tgm3, Serpinb3a, Lor*) (**Figure 7H-I**). The first differentiating cells have been reported from approximately E13 (Damen et al., 2021; Lechler and Fuchs, 2005), and in line with this we also we also detect the first *EPI EarlyDiff* cells at E13.5. However, mature *EPI LateDiff* cells do not appear until E14.5 (**Figure S6C**). *In situ* staining revealed that some rare *Krt5*^+^ basal layer cells already start upregulating *Krt10* and *Krtdap* even though they haven’t delaminated yet (**Figure 7J**). Interestingly, in embryonic skin the suprabasal expression pattern of *Krt5* differs from that of *Krt14*, even though those two keratins usually co-polymerize. *Krt14* is strictly downregulated in suprabasal cells, while *Krt5* is still expressed in suprabasal cells – albeit at slightly reduced levels (**Figure 7I** and **S6G**). The only other embryonic epidermal population where we found expression of those two keratins differing is the hair placode, where *Krt5* but not *Krt14* is downregulated (**Figure S6H**).

As the early signals that make a basal cell commit to differentiation are not fully resolved, we utilized our dataset to identify potential drivers. In order to reveal the differentiation trajectory, we had to perform rigorous cell cycle corrections, as this was the dominating factor in initial velocity analysis. The corrected data revealed a clear differentiation trajectory from *EPI Basal2/4* cells toward *EPI EarlyDiff/LateDiff* cells, that we could use to model a pseudotime reflecting differentiation (**Figure S6I**). We then filtered for the pseudotime-dependent genes, that come up very early in pseudotime (**Figure 7K**). *Notch1*, a known commitment switch in epidermal differentiation (Blanpain et al., 2006; Rangarajan et al., 2001), *Cdh1* (better known as E-CAD), which is responsible for altered adhesion properties that allow keratinocytes to differentiate (Miroshnikova et al., 2018), and *Grhl3*, which facilitates epidermal stem cell differentiation (Lin et al., 2020) were among the top hits in our list suggesting that this list may include unknown differentiation drivers. Furthermore, the upregulation of these genes seemed to be transient or at least most pronounced in early differentiating cells supporting a potential switch-like function (**Figure 7K** and **S6I**).

### Hair placodes engage in the establishment of blood vessels, nerves, and immune environment

The epithelial counterpart to the dermal condensate, which is necessary for hair follicle formation, is the hair follicle placode. While placodes became morphologically first visible at E14.5 (**Figure S1H,I**), based on mRNA expression we already identified at E13.5 the cells that prepare for a placode fate (*EPI EarlyPlacode* and *EPI LatePlacode*) (**Figure 7A,C** and **S6A,C**). These cells show intermediate expression levels of typical placode markers such as *Fgf20* and *Dkk4* but lack detectable expression of more mature markers like *Shh* and *Lhx2* (**Figure 7L**) (Levy et al., 2005; Rhee et al., 2006). By staining for *Dkk4* and *Ltb* mRNA, we could even capture a few of those early placode cells *in situ* in E13.5 epidermis (**Figure 7M-N**). *Shh* mRNA could be stained in more mature placodes at E14.5 and always located to the very center of the placode (and to the very tip of the UMAP) (as shown in Morita et al., 2021), while other markers such as *Ltb* showed a slightly broader expression pattern (**Figure 7O-P** and **S6J**). Interestingly, while placode cells upregulated numerous markers, they only downregulated a handful of genes such as *Gas1*, *Krt5*, and *Hspb1*. It thus seems likely that placode commitment is rather determined by the gain than the loss of expression (**Figure 7L**). While reporter mice and *in situ* mRNA stainings have long revealed that placode patterning begins prior to E14.5 (Ahtiainen et al., 2014; Fliniaux et al., 2008; Närhi et al., 2008; Zhang et al., 2009), previous scRNA-seq studies of embryonic skin did not reveal those E13.5 cells with early placode markers likely due to the choice of a different analysis strategy (Gupta et al., 2019).

Finally, our receptor-ligand analysis (**Figure 6A-C**) suggests that the just formed placode and dermal condensate cells immediately engage in a number of reciprocal interactions with several major cell types known to be crucial for proper hair follicle function. The mature hair follicle depends on blood supply, innervation, and support from tissue resident immune cells (Adachi et al., 2015; Brownell et al., 2011; Wang et al., 2019; Xiao et al., 2013). Through expression of key factors, the earliest stage of developing hair follicles seems to ensure that the newly formed hair follicle is properly innervated (via neurotrophic factors such as *Bdnf, Nrtn, Ntf3*, and *Edn3*), surrounded by blood vessels (via angiogenic factors such as *Bmp2, Bmp7*, and *Mfge8*) and supported by immune cells (via immune-recruiting and activating factors such as *Tnf* and *Ltb*) (**Figure S6K**).

## DISCUSSION

This work presents an unprecedented picture of early skin development. Through comprehensive computational analysis of all cell types sampled at E12.5, E13.5, E14.5, cell-type localization *in situ,* and *in vivo* cell-fate mapping, we answered major outstanding questions in mouse skin development and made unexpected new discoveries. When and where does skin begin? How heterogeneous are fibroblasts prior to hair follicle placode formation? Is the periderm merely a signaling-inert protective layer to be shed at birth?

### When and where does skin begin – setting new anatomical and molecular landmarks

Until now, E12.5 dermis and non-skin-associated cells were perceived as a seemingly homogenous tissue covering the area between the vertebrae and epidermis. Similarly, little was known about dermal tissue architecture and cell type diversity at E13.5 and 14.5. Thus, it was critical to unbiasedly sample the skin and the underlying tissue at full thickness from E12.5-14.5 (see **Methods**) in order to define the developing “skin-associated” tissue space. In this work, we iteratively established anatomical and molecular tissue landmarks of the skin and underlying tissue.

One of the most important landmarks to define cell populations as skin-associated in mouse is the PCM. While it is well-known that the PCM originates from the dermomyotome, most specifically from the E9.5 Pax7+ lineage (Amini-Nik et al., 2011; Atit et al., 2006; Lepper et al., 2011), its histological emergence had not been documented. Through histological analysis of the developing muscle structures we defined the emergence of the PCM (**Figure 1D-F**). Subsequent smFISH stainings (using muscles layers and cell-membranes as a landmark) enabled us to place our newly defined scRNA-seq subpopulations in the tissue context. Altogether, this resulted in a detailed molecular tissue guide that complements previous findings and accelerates the interpretation of future findings.

### Fibroblast heterogeneity and the emergence of papillary and reticular dermis

It has been an accepted view that at E12.5, E13.5, and E14.5 dermal fibroblasts constitute a ‘uniform cell type’ where each cell can still differentiate into all fibroblasts of postnatal skin, and that it is not before E16.5 when dermal fibroblasts commit to two different lineages which generate either the upper dermis or the lower dermis (Driskell et al., 2013). Our data reveals that molecular and functional diversity of fibroblasts is already established at E13.5, with further fibroblast specifications happening at E14.5 when the dermal condensate forms (**Figure 3**). We also find clear transcriptional heterogeneity in E12.5 dermis (**Figure 2**), which based on calculating the likelihood of fate outcomes *in silico* suggests a fate bias towards distinct future fibroblast subtypes. To what degree these E12.5-14.5 lineages remain plastic or are already fate-restricted under homeostatic conditions remains to be determined.

Driskell et al. also established a molecular distinction of fibroblasts into papillary dermis (DPP4^+^/DLK1^-^/LY6A^-^), reticular dermis (DPP4^-^/DLK1^+^/LY6A^+^) and hypodermis (DPP4^-^/DLK1^-^/LY6A^+^) starting from late embryogenesis. They detected DLK1 protein expression throughout the dermis until E16.5, while lineage-specific DPP4 (also known as CD26) and LY6A emerged around E16.5 (Driskell et al., 2013). In our data, *Dlk1* was similarly expressed throughout the dermis. Additionally, we detected cells with a *Ly6a*^+^/*Dpp4*^+^ double signature in the *FIB Inter* population starting from E13.5 (**Figure S3F**), raising the question if there is a relationship between our *Dpp4*^+^/*Ly6a*^+^ *FIB Inter* cells and the papillary and reticular dermis. As we observe *FIB Inter* cells at a time when *FIB Upper* and *FIB Lower* cells (the tentative precursors of papillary and reticular dermis; *Dpp4*^-^/*Ly6a*^+^) have already been established, it is likely that *FIB Upper* and *FIB Lower* cells acquire *Dpp4* or *Ly6a* expression independently of *FIB Inter* cells.

### Revisiting early progenitors of dermal and subcutaneous white adipose tissue

In 2013, Wojciechowicz et al. postulated a possible presence of pre-adipocytes already at E14.5, which our scRNA-seq data clearly supports (Wojciechowicz et al., 2013). Cells within the *FIB Inter2/3* populations increasingly express typical adipogenic genes, such as *Pparg* and *Cebpa* (increasing expression from *FIB Inter2* to *FIB Inter3*), suggesting the presence of adipocyte precursurs (**Figure 3I** and **S3H**). *In situ* staining for *Pparg* mRNA and protein in E14.5 skin (**Figure S3I**) revealed *Pparg*^+^ cells within the subcutanous interstitium, where the subcutanous white adipose tissue (SWAT) will form, as well as a few *Pparg*^+^ cells just above the PCM, where the dermal white adipose tissue (DWAT) will form (Driskell et al., 2014). Notably, a recent study found that *Dpp4*^+^*/Anxa3*^+^*/Wnt2*^+^ fibroblasts – a description fitting our *FIB Inter2/3* cells – can function as adipose mesenchymal progenitors in human as well as murine skin (Merrick et al., 2019). Moreover, following a recently suggested nomenclature (Ferrero et al., 2020), *Dpp4* expression combined with positivity for *Ly6a* and *Cd55* would classify our *FIB Inter2/3* cells as adipose stem cells (ASCs) (**Figure S3F-G**).

Given this concurrence with the literature, it was very surprising that we did not observe any Gata6-Tom traced adipocytes. As we started tracing at E13.5, when the SWAT and DWAT are not yet separated by the PCM, we in principle could have found Tom-traced cells in both adipose compartments, but we certainly expected Tom-traced adipocytes in one of them. However, we did not find traced DWAT cells and due to technical limitations SWAT was lost when harvesting postnatal skin. This leaves us with three open possibilities: i) *FIB Inter* cells do not represent adipocyte precursors at all, which is less likely based on the expression of typical adipogenic genes, ii) tracing at E13.5 is not efficient enough to label the adipose progenitors but tracing at E14.5 might have been, or iii) *FIB Inter* cells only contribute to SWAT formation and thus DWAT and SWAT originate from independent precursors, which seems more likely based on its location on opposite sides of the developing PCM and supported by the current view that DWAT is morphologically and developmentally distinct from SWAT (Driskell et al., 2014; Wojciechowicz et al., 2013). As the latter view is derived from experiments performed from E14.5 onward, the earliest determination of a fibroblast subset towards generating adipose tissue remains an interesting route to be explored.

### Unexpected heterogeneity in E12.5 epidermis and a surprisingly signaling-rich periderm

We identify significant heterogeneity in E12.5 epidermis, that to date had been considered as a uniform epithelial sheet. However, the function of the newly identified, highly distinct *EPI BasalTagln* population remains elusive. The abundance of receptors and ligands suggests a possible role for providing transient signaling during skin development (e.g. to activate the upper dermis via Wnts) (Collins et al., 2011; Donati et al., 2014; Lichtenberger et al., 2016). However, the *EPI BasalTagln* population could also be a cell source for the periderm, since a number of genes (at lower levels) overlap with the periderm signature (**Table S1**). In comparison, the remaining IFE basal cells at E13.5 and 14.5 were transcriptionally unexpectedly similar. As we did not identify any cluster-specific gene expression in *Basal2-4*, this separation is unlikely to represent cell populations with distinct behavior or function, but rather cell cycle influences. Moreover, we made the interesting observation of rare differentiating basal cells (**Figure 7J**), reminiscent of delaminating K10+ cells in adult mouse skin (Cockburn et al., 2021). It is tempting to speculate that epidermal stratification of embryonic skin may be fueled by two coexisting basal-cell behaviors: i) delamination triggered by basal cell crowding (predominant mechanism) (Damen et al., 2021; Miroshnikova et al., 2018), and ii) delamination through gradual differentiation which is the main mechanism in postnatal and adult skin (Cockburn et al., 2021; Lin et al., 2020).

The existence of periderm cells in embryonic skin has long been known (M’Boneko and Merker, 1988). However, periderm cells did not form their own cell cluster in any of the previous scRNA-seq studies likely due to low cell numbers (Fan et al., 2018; Ge et al., 2020; Gupta et al., 2019). As periderm cells also express typical IFE differentiation genes such as *Grhl3* and *Zfp750* (**Figure 7F,H**) there is a risk of misinterpreting periderm cells as differentiating cells and our characterization of the periderm signature may help the interpretation of future studies. Notably, the periderm not only proliferates (Richardson et al., 2014), indicating the potential of self-maintenance until it’s shed, but it is also highly signaling prone (**Figure S6E**) which raises the possibility of previously unrecognized “non canonical” periderm functions in embryonic skin development.

### Limitations and advances

The most insightful transcriptional investigations of early embryonic skin to date relied on known markers and averaged transcriptomes (bulk RNA-seq of FACS-sorted populations) (Sennett et al., 2015) or focused their scRNA-analysis on specific cell types/processes such as the molecular origin of IFE cells (Fan et al., 2018), the cellular origin of hair follicle stem cells (Morita et al., 2021), or placode- or dermal condensate-fate specification (Gupta et al., 2019; Mok et al., 2019).

In this work we combined in-depth analysis of all randomly sampled cells present in the scRNA-seq dataset with spatial tissue mapping to generate the most detailed molecular and spatial picture of embryonic skin development today. Due to the vast amount of data, we had to focus our detailed analysis on a number of major outstanding questions, such as the emergence of skin and early fibroblast heterogeneity. However, we would like to acknowledge that several of the scRNA-seq results at the current state may represent data-driven suggestions rather than final conclusions, and future studies focusing on specific aspects will be necessary to put these findings into the bigger context. Overall, we are convinced that this and future studies that present a holistic picture of cell types – which together orchestrate organ function – are of very high value as they advance our understanding from studying individual cell types (in isolation) to their communal functions at the tissue level.

## Contribution

M.K. and T.J. formulated the research question and designed the study. T.J. performed all single-cell sequencing experiments including animal work and cell isolation. T.J. and T.D. sampled tissue for validation experiments. T.J. performed the majority of bioinformatic data analysis. K.A., P.C., and Å.B. contributed to bioinformatic data analysis. T.J. planned and set up validation stainings (protein-IF and RNA-FISH). T.J. and K.A. performed and imaged validation stainings. T.J. and C.L.L. performed lineage tracings in mice and K.A. and M.K. analyzed the tracing patterns. T.J. and K.A. performed image processing. B.M.L., M.R., M.L.M., T.D., M.E.K. helped to interpret the results. G.D. provided Gata6-tracing mice. M.K. and T.J. wrote the manuscript with input from all authors.

## Acknowledgements

We would like to thank Adelheid Elbe-Buerger for valuable scientific discussions; Hong Qian and Lakshmi Sandhow for generously sharing Ebf2-tracing mice; Thibault Bouderlique for sharing his expertise on embryo dissections; Stefania Giacomello (from National Bioinformatics Infrastructure Sweden at SciLifeLab), Simon Joost, Christoph Ziegenhain, and Rickard Sandberg for advice on computational analysis; Alexandra Are and Xiaoyan Sun for help with mice and stainings; and Makbule Sagici for embedding and cutting paraffin samples. This work was supported by grants from the Swedish Research Council (VR), Swedish Foundation for Strategic Research (SSF), Center for Innovative Medicine (CIMED), Swedish Cancer Society (Cancerfonden), Karolinska Institutet, and Ragnar Söderberg Foundation to MK, Karolinska Institutet KID funding to TJ and KA, and a Wenner-Gren postdoctoral fellowship to TD. The G.D. laboratory is supported by AIRC (MFAG 2018 ID: 21640) and a Chan Zuckerberg Initiative Grant (DAF2020-217532). P.C. and Å.B. are financially supported by the Knut and Alice Wallenberg Foundation as part of the National Bioinformatics Infrastructure Sweden at SciLifeLab. All single-cell experiments were performed at the Eukaryotic Single Cell Genomics core facility, SciLifeLab, Stockholm. The authors also acknowledge support from the National Genomics Infrastructure in Stockholm funded by Science for Life Laboratory, the Knut and Alice Wallenberg Foundation and the Swedish Research Council, and SNIC/Uppsala Multidisciplinary Center for Advanced Computational Science for assistance with massively parallel sequencing and access to the UPPMAX computational infrastructure. Imaging was performed at the LCI facility/Nikon Center of Excellence, Karolinska Institutet, supported by grants from the Knut and Alice Wallenberg Foundation, Swedish Research Council, KI infrastructure, Centre for Innovative Medicine and Jonasson center at the Royal Institute of Technology.

## STAR METHODS

### KEY RESOURCES TABLE

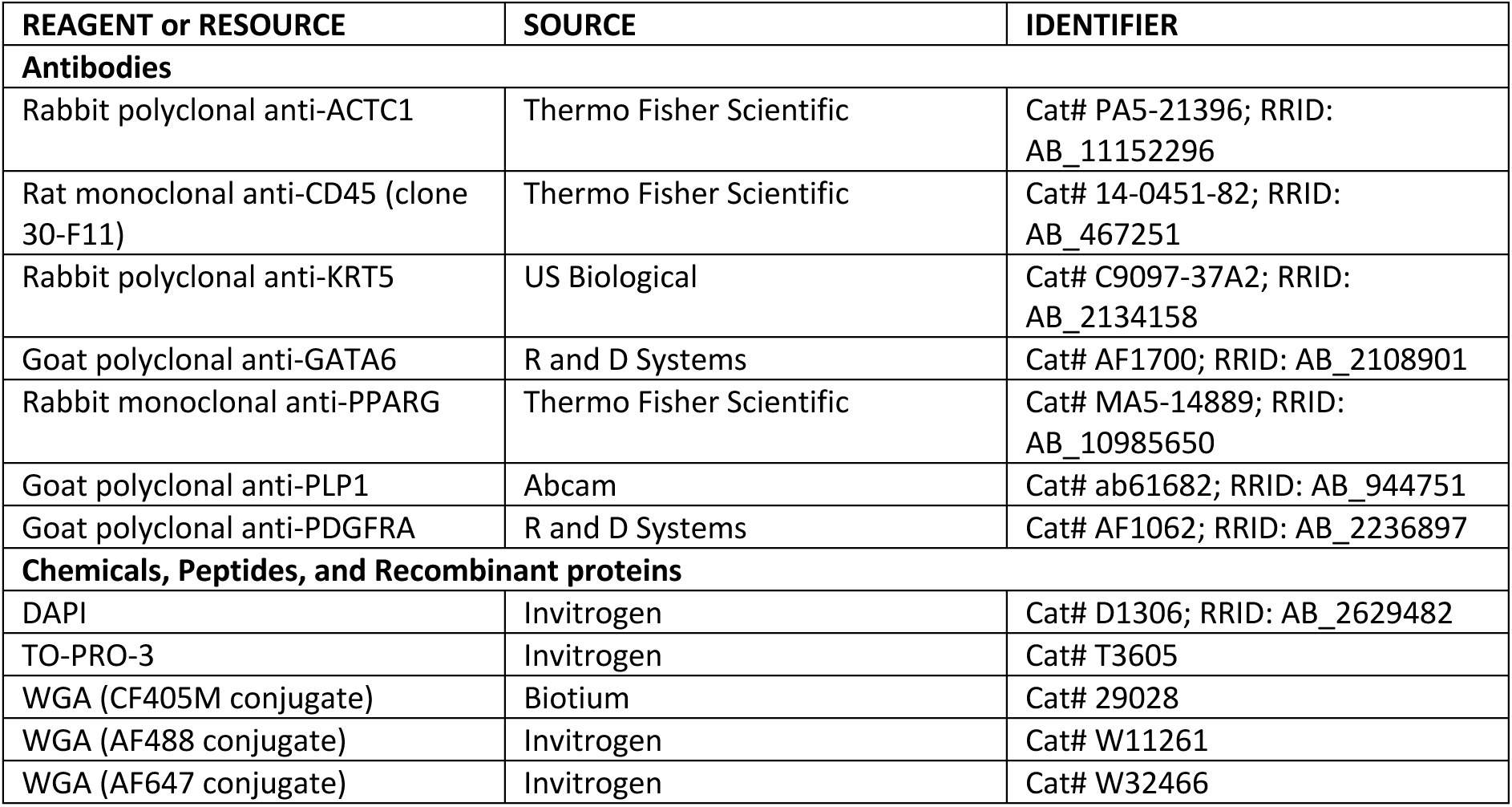
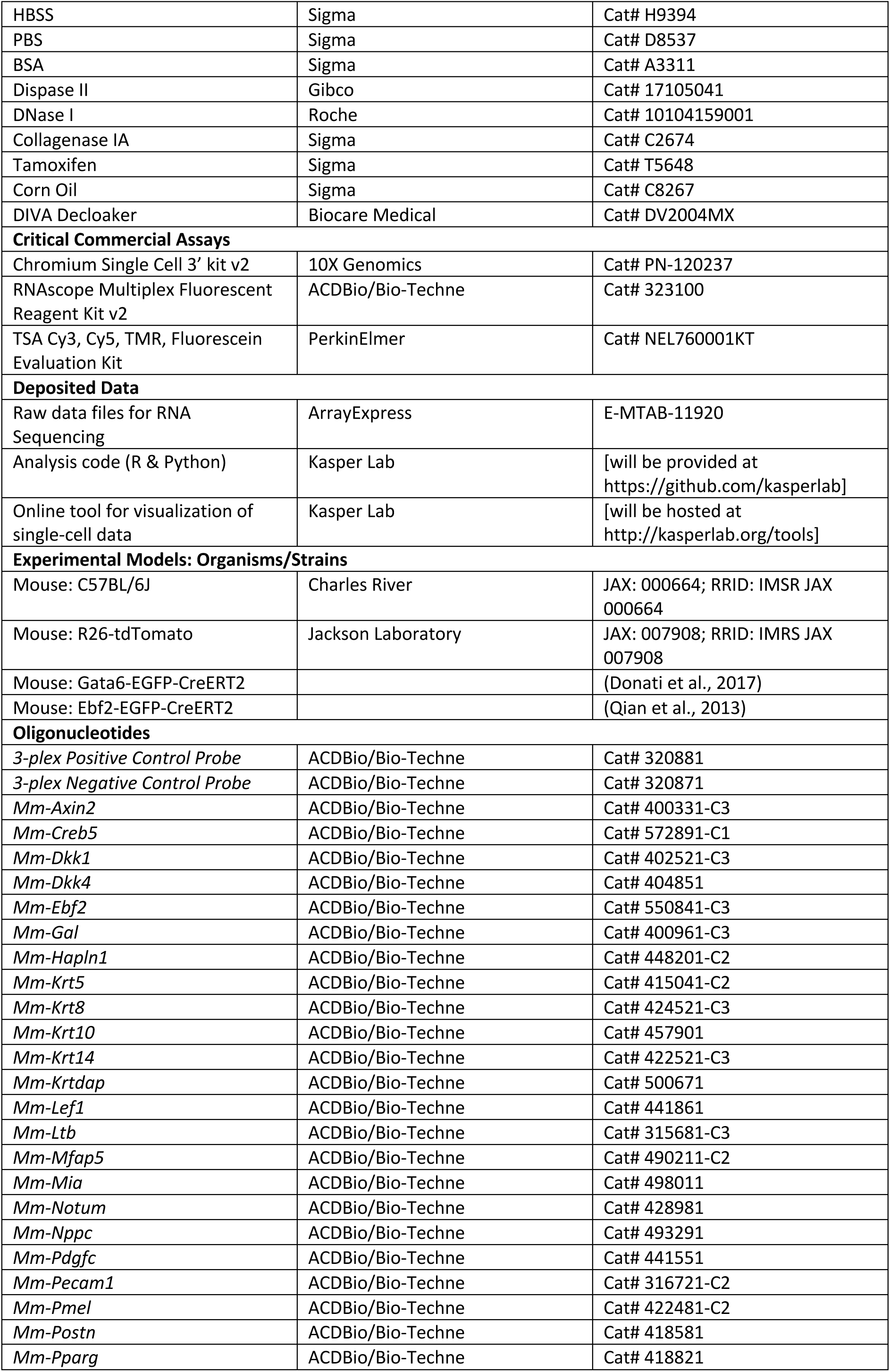
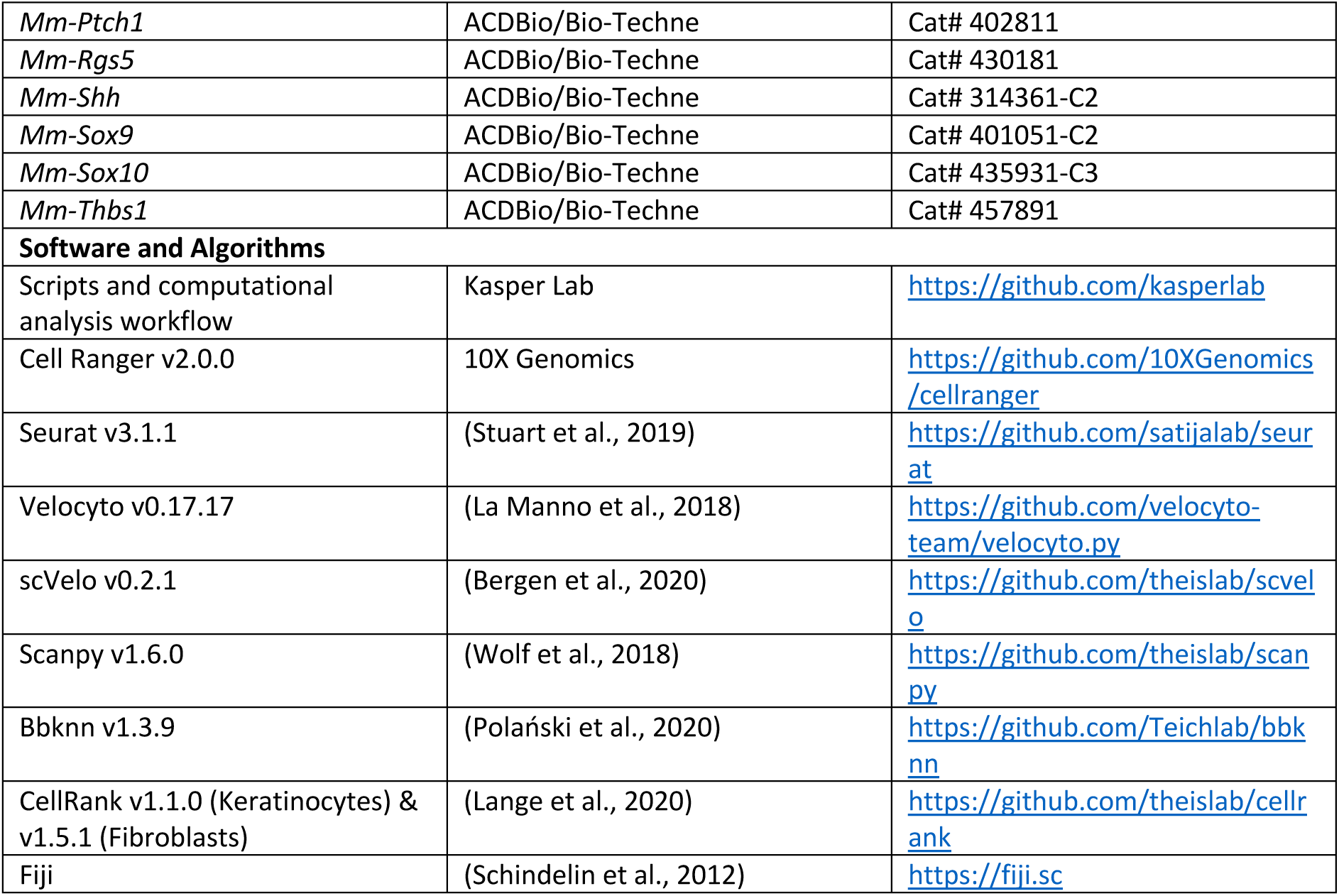

### LEAD CONTACT AND MATERIALS AVAILABILITY

Further information and requests for resources and data should be directed to and will be fulfilled by the lead contact Maria Kasper. This study did not generate new unique reagents.

### EXPERIMENTAL MODEL AND SUBJECT DETAILS

#### Mouse work

The study was performed on wild-type C57BL/6J mouse embryos (mix of males and females - gender was only determined in retrospect from the sequencing data (**Figure S1E**)). Timed matings to obtain embryos of specific embryonic ages were set up in the evenings and the next morning was defined as E0.5. Pregnancy after timed matings was determined by comparing weight difference between the start of the mating and 10 days after. Pregnant moms were sacrificed by cervical dislocation when embryos reached the embryonic age of 12.5, 13.5, or 14.5 days, respectively, and embryos were processed for cell isolation or paraffin-embedding. Lineage-tracing experiments were performed by crossing previously described Gata6-EGFP-CreER^T2^ (Donati et al., 2017), Ebf2-EGFP-CreERT2 (Qian et al., 2013) and *R26-tdTomato* knock-in strains (Madisen et al., 2010) (hereafter Gata6-Tom). Gata6-Tom mice received i.p. injection of 2mg tamoxifen (in corn oil at a concentration of 20mg/ml) at embryonic day 13.5. Uninjected mice were used as leakiness control. Tissues were sampled either 2 days after induction of lineage tracing (i.e., E15.5) or postnatally (postnatal day 5). Ebf2/Tom mice were i.p. injected with 2mg tamoxifen (in corn oil at a concentration of 20mg/ml) at E14.5 and tissues were sampled either 2 days after induction of lineage tracing or at E18.5.

FELASA recommendations for harmonized health monitoring were followed. The mice were fed *ad libitum* and handled and housed under standard conditions. All mouse work (except Gata6-lineage tracings) was performed in the animal facility of Karolinska University Hospital Huddinge and in accordance with Swedish legislation and approved by the Linköping Animal Ethics Committee. Gata6-lineage tracings were performed in the animal facility of the Molecular Biotechnology Center at the University of Turin and in accordance with Italian legislation and approved by the local Animal Ethics Committee and the Italian Ministry of Health.

### METHOD DETAILS

#### Replicates

Sequencing was performed on five embryos per embryonic time point. These five embryos originated from two litters and were sampled on two different days. All 15 samples were processed and sequenced individually and can thus serve as true biological replicates (**Table S2**).

Each individual staining was performed on skin samples from at least 3 different embryos per embryonic age.

#### Tissue embedding

Whole embryos and postnatal skin tissue were formaldehyde-fixed in 4% PFA for 24h at room temperature and subsequently processed for FFPE sections (4um thickness). When sectioning whole embryos, tissue sections were collected close to the dorsal midline.

#### Fluorescent *in situ* hybridization (FISH)

For independent validation and mapping of cell populations, single-molecule FISH was performed using the RNAscope Multiplex Fluorescent Detection Kit v2 according to manufacturer’s instructions using TSA with Cy3, Cy5, and/or Fluorescein on FFPE sections of the embryos. The used probes are listed in the Key Resources Table. All sections were counterstained with either WGA-405 (1:50), WGA-488 (1:50), WGA-647 (1:50), DAPI (1:500), TO-PRO3 (1:1000) or combinations of them.

#### Immunofluorescence (IF)

Immunofluorescence was performed either alone or after completed RNAscope staining. Combined with RNAscope, sections were washed in TBST once and then blocked and stained as in regular IF stainings. For IF without RNAscope, antigen retrieval was performed using DIVA Decloaker. The following antibody concentrations were used: ACTC1 (1:500), CD45 (1:200), KRT5 (1:50), GATA6 (1:25), PPARG (1:100), PLP1 (1:1000), and RFP (endogenous tdTomato is lost during FFPE processing, 1:200). All sections were counterstained with either WGA-405nm (1:50), WGA-488nm (1:50), WGA-647nm (1:50), DAPI (1:500), TO-PRO3 (1:1000) or combinations of them.

#### Imaging and image analysis

Images were acquired on a Nikon A1R spinning disk confocal as tiled images (10%–15% overlap) and stitched by NIS Elements software. Subsequently, all images were processed in a uniform way (maximum intensity projection, background removal with the ‘‘subtract background’’ plug-in, brightness adjustment, pseudo-colouring) using Fiji (Schindelin et al., 2012).

#### Cell isolation

Dorsal skin of embryos (**Figure S1B**) was dissected with the help of fine dissection tools and dissected skins were incubated in Dispase II (2mg/ml), Collagenase IA (0.2%), and DNAse I (20U/ul) in PBS for 40 minutes at 37°C in ultra-low attachment plates (Corning Costar) on an orbital shaker. The obtained cell suspension was passed through a 40 mm cell strainer. The flow-through was spun down, and subsequently resuspended in PBS + 0.04% BSA. Samples were transported to core facility in PBS + 0.04% BSA in Eppendorf tubes that had been coated with PBS + 20% BSA overnight. Viability of the cell suspension was determined using trypan blue on an EVE automatic cell counter.

Of note, when peeling off dorsal skin tissue there were no means to technically prevent co-isolation of cells from the tissue layers underlying skin (cells from the deeper muscle layers, interstitial cells, and/or chondrocytes) Biologically, sampling the entire embryonic outer layer (skin and underlying tissue) was important as those embryonic timepoints are/were ill-defined in terms of what can be considered skin tissue.

#### Library preparation, sequencing and processing of sequencing data

Single-cell cDNA libraries were prepared using the 10X Genomics Chromium Single Cell 3’ kit v2 according to the manufacturer’s instruction. Libraries were sequenced on the HiSeq2500 system (Illumina). Raw sequencing data was processed using the 10X Genomics Cell Ranger package and the mm10 reference genome.

### QUANTIFICATION AND STATISTICAL ANALYSIS

#### Data analysis

##### Analysis workflow

All downstream data analysis was performed using a mix of custom scripts and published analysis packages as described below and in **Figure S1F**, utilizing a mix of R packages (most importantly Seurat) as well as Python packages (most importantly Scanpy and scVelo) (Bergen et al., 2020; Stuart et al., 2019; Wolf et al., 2018).

Major decisions on analytical approaches will be presented below, while we refer to the pipelines that will be deposited on GitHub (https://github.com/kasperlab) for any questions regarding details such as chosen parameters.

##### Quality control and pre-processing

Cell-filtering was performed by sample and was based on the following criteria: (a) remove cells with <200 genes/cell, (b) remove cells with low diversity index, i.e. Shannon and inverse Simpson index (this removes red blood cells, that are naturally expressing only a small variety of genes), and (c) remove cells that are simultaneously in the lowest 0.05% quantile for genes/cell (nFeature) and for reads/cell (nUMI) and that have a contribution of mitochondrial genes of <1% or >10%. By using these combinatorial criteria, it was ensured that cells would not be excluded just because they have e.g. a lower respiratory rate (i.e., low mitochondrial percentage only).

Subsequently, all 15 samples were combined into one full dataset and filtered once more on genes being expressed in at least 5 cells. Ribosomal genes (Rps and Rpl gene families), haemoglobin genes (Hba and Hbb gene families), as well as mitochondrial genes (mt gene family) were removed, as they interfered with the identification of meaningful marker genes. Log-normalization was performed using Seurat’s NormalizeData function.

##### Determining sex of embryos

As it was very challenging to determine the sex of the embryos during sampling (due to early developmental stage and the need to process samples quickly for sequencing), litter mates were randomly chosen for sequencing and their sex was determined in retrospect from the scRNA-seq data based on the percentage of reads coming from the X chromosome and the Y chromosome (**Figure S1E**). This data revealed the gender identity for each of the embryos (3 females/2 males for E12.5; 1 female/4 males for E13.5; 1 female/4 males for E14.5).

##### Removal of cell doublets and low-quality cells

During analysis, two small groups of doublets (keratinocyte-fibroblast doublets that clustered with keratinocytes and pericyte-fibroblast doublets that clustered with fibroblasts) were encountered as well as some low-quality keratinocytes that survived global quality control (low nFeature, low nUMI, low perc.mito). Those cells were removed, and analysis was re-run without them.

Furthermore, one cluster was identified during first-level clustering which very likely corresponds to neuronal cells (sensory neurons defined by e.g. *Neurod1* and *Pou4f1* as well as sympathetic neurons defined by e.g. *Stmn2* and *Nefm*). While the signature was rather clean, the cell population originated only from a single E13.5 embryo and thus was not reproducible and most likely the result of some tissue sampling issue. Hence, the cluster was removed. Please refer to the results section, for a discussion of why neuronal transcriptomes would not be expected in this dataset.

##### Removing effect of confounding factors

To counteract a slight batch effect (i.e., slightly differing characteristics such as higher percentage of histone reads and pseudogene reads) linked to one of the sampling days, linear regression was performed using Seurat’s ScaleData function – a rather mild measure for data integration. Regression was performed for sampling date, as well as gender, percentage of mitochondrial genes, total read counts, and cell cycle scores (S.Score and G2M.Score) as those could also potentially influence dimensionality reduction and clustering while not representing the biological variables of interest.

##### Prediction of cell cycle stage

Cell cycle stage was predicted using Seurat’s CellCycleScoring function.

##### Generation of loom files

To allow for running RNA velocity analysis on spliced and unspliced mRNAs, we generated loom files using Velocyto’s run10x function with default parameters and using the mm10 reference genome.

##### Feature selection

Feature selection was performed using the mean-dropout-method originally suggested by (Andrews and Hemberg, 2019) in our own implementation. The 3000 genes with the highest dropout rate given their mean expression level (across non-zero counts) were chosen to be included in further analysis.

Feature selection was performed separately for the full dataset, fibroblasts, or keratinocytes, respectively, to allow for the detection of more subtle differences within fibroblasts and keratinocytes, respectively, that were hidden in the full dataset where distinct signatures of major cell types dominate the highly variable genes.

##### Clustering, spatial embedding, and trajectory analysis

Clustering, spatial embedding and trajectory analysis were separately adjusted for each of the three analysed groups (full dataset, fibroblasts, and keratinocytes) as they possessed very dissimilar features. The full dataset contained very distinct cell types, while fibroblasts and keratinocytes constituted a much more homogenous cell population with more gradual expression changes. Also, the biological questions that were of interest differed strongly, so different aspects had to be emphasized and analysis was adjusted accordingly.

##### Full dataset

Dimensionality reduction was performed in Seurat using PCA with the most highly variable genes as input after initial scaling with Seurat’s NormalizeData function (a scaling factor of 10 000 was chosen as this roughly reflects the median reads/cell among the filtered cells). Subsequently, hierarchical clustering (hclust function) was performed based on PCA-reduced data. While this clustering worked well without any further need for data integration, the downstream dimensionality reduction (UMAP) still showed signs of sampling date-derived batch effects. Thus, a batch-corrected neighbourhood graph from BBKNN (Polański et al., 2020) was used to prevent batch-derived separation in UMAP space. To this aim, the regressed dataset was transferred to Python, principal components were recalculated, BBKNN was run, and dimensionality reduction was performed using Scanpy’s UMAP function, which was then used to display the results of the hierarchical clustering (**Figure 1B**).

##### Fibroblasts

Fibroblast batch correction was similarly done in Scanpy using BBKNN for UMAP representation. Next, cell clustering was performed using the Leiden algorithm (Traag et al., 2019), revealing the fibroblast subpopulations for subsequent analysis (**Figure S2A**). As the dermal condensate (DC) is a structure of great importance to skin development, we decided to further subcluster the DC into *FIB EarlyDC* and *FIB LateDC* using the Leiden algorithm.

Next, we imported spliced and unspliced mRNA information (from loom files) for RNA-velocity analysis using scVelo’s velocity function in the stochastic mode on the highly variable genes. The predicted dynamics were then plotted on top of the pre-computed UMAP (**Figure S2B**).

To further analyse cellular dynamics towards the endpoints, we used CellRank (v1.5.1) pseudotime kernel. As input to the model, we calculated velocity pseudotime with identified root cells in the FIB origin populations and end cells as extreme points based on diffusion maps. Finally, we calculated absorption probabilities for each cell to become any of the identified end points.

##### Keratinocytes

Dimensionality reduction was performed in Seurat using PCA with the most highly variable genes as input after initial scaling. Subsequently, hierarchical clustering (hclust function) was performed based on UMAP-reduced data. The regressed keratinocyte dataset was then transferred to Python and combined with the loom-file-derived information on spliced and unspliced mRNAs.

To better understand epidermal stratification, a subset of E14.5 cells was studied, as E14.5 is the first embryonic age to capture the full differentiation trajectory. Cells from the *EPI Basal1*, *EPI Basal2*, *EPI Basal3*, *EPI Basal4*, *EPI EarlyDiff*, and *EPI LateDiff* clusters were included in the analysis. Cells related to placode, periderm, or the *EPI BasalTagln* cluster were excluded as they could potentially interfere with the differentiation trajectory. The subset was processed as described before (PCA, BBKNN, UMAP) and velocity analysis was performed, which revealed striking dominance of the cell cycle in RNA-velocity predictions (**Figure S6I**). Thus, cell cycle effects were regressed out in spliced and unspliced mRNA using Scanpy’s regress_out function, which resulted in a striking velocity pattern reflecting epidermal stratification (**Figure S6I**). Finally, CellRank in combination with diffusion maps and RNA-velocity based pseudotimes was used to find macrostates and generate a refined pseudotime, to calculate lineage drivers, and to fit a GAM model for gene expression analysis (Lange et al., 2020). The top 200 pseudotime-dependent genes were obtained and the 40 genes among them with the earliest peak in pseudotime were displayed (**Figure 7K**).

Using a similar approach, the differentiation trajectory within the *EPI Periderm* cluster was analysed. Periderm cells from E14.5 were identified as a terminal state and a periderm maturation trajectory could be modelled. The top 200 pseudotime-dependent genes were obtained and the 13 genes among them with the latest peak in pseudotime were displayed (**Figure S6F**).

##### Test for differential expression of genes in cell populations

Marker genes overexpressed in certain cell populations were determined using the Wilcoxon rank sum test in the Seurat implementation. Marker genes were required to be detected in at least 20% of cells in the respective population and to have a (natural) log fold change ≥ 0.25 compared to all other cells. Correction for multiple testing was performed using the Bonferroni method and the threshold for the FDR (false discovery rate) adjusted p-value was set to 0.05 (**Table S1**).

##### Receptor-ligand interactions

Receptor-ligand pairing was based on the approach presented by (Joost et al., 2018). In brief, receptors and ligands contained in the marker gene list were considered for potential receptor-ligand pairs. For each cluster pair, receptor-ligand interactions were identified by querying a receptor-ligand database (**Figure S5**). In contrast to (Joost et al., 2018), the curated receptor-ligand databases from (Ramilowski et al., 2015) and (Cabello-Aguilar et al., 2020) were combined to obtain an even more complete set of potential interactions. Both databases are based on human data, but we assume that the majority of registered interaction pairs are also valid for homologous genes/proteins in mice. The code was furthermore optimized for run-time and parallel computing.

To test for the enrichment of receptor-ligand pairs between two populations, the observed number of receptor-ligand pairs was compared to the number of pairs obtained from an equally sized randomly sampled pool of receptors and ligands. For each cluster pair, this simulation was repeated 10 000 times and significantly enriched interactions (p ≤ 0.05 for Benjamini-Hochberg-corrected p-values) were combined into **Table S3**. Each ligand and receptor in this table was manually annotated with reported functions and each pair was manually scored for likely involvement in the development of vessels, nerves and the immune system (**Figure 6, Table S3**).

Receptor-ligand interactions were identified within and between major cell populations, fibroblast subpopulations, and epidermal subpopulations (any possible combination of them with any possible signalling directionality).

### DATA AND CODE AVAILABILITY

#### Data resources

The sequencing data reported in this paper will be deposited in GEO.

#### Software

The complete computational analysis workflow as well as a list with the complete cluster assignment will be available at https://github.com/kasperlab.

### DATA PORTAL

Accompanying this work, upon publication we will provide a user-friendly data portal for exploring the single-cell data, similar to our previous publications (http://kasperlab.org/tools).

