## Supplemental Figures 1-6 for "Molecular and spatial design of early skin development"

**Figure S1.** Details of experimental approach and quality control, Related to Figure 1.

**Figure S2.** Deconstruction of fibroblast heterogeneity at E12.5 (expression and location), Related to Figure 2.

**Figure S3.** Deconstruction of fibroblast heterogeneity at E13.5 and E14.5 (expression and location), Related to Figure 3.

**Figure S4.** Cell types contributing to embryonic skin besides fibroblasts and keratinocytes, Related to Figure 5.

**Figure S5.** Workflow for receptor-ligand interactions, Related to Figure 6.

**Figure S6.** Epidermal development from a single basal layer towards a HF-inducing and stratified epithelium, Related to Figure 7.

### Figure S1

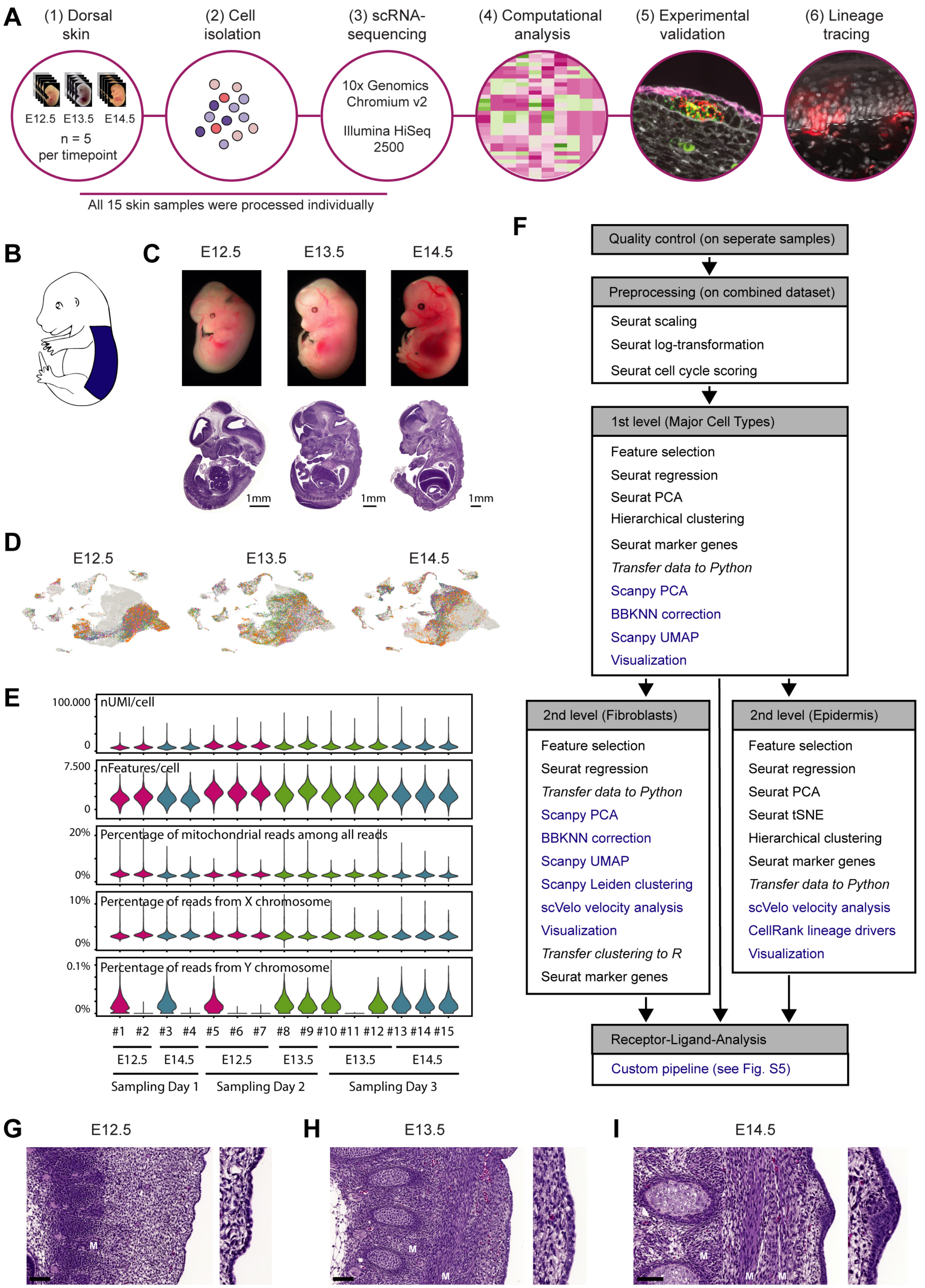
A
(1) Dorsal skin
(2) Cell isolation
(3) scRNA-sequencing
(4) Computational analysis
(5) Experimental validation
(6) Lineage tracing
E12.5 E13.5 E14.5
n = 5 per timepoint
10x Genomics Chromium v2
Illumina HiSeq 2500
All 15 skin samples were processed individually
B
C
E12.5 E13.5 E14.5
1mm 1mm 1mm
D
E12.5 E13.5 E14.5
E
100.000 nUMI/cell 0
7.500 nFeatures/cell 0
20% Percentage of mitochondrial reads among all reads 0%
10% Percentage of reads from X chromosome 0%
0.1% Percentage of reads from Y chromosome 0%
#1 #2 #3 #4 #5 #6 #7 #8 #9 #10 #11 #12 #13 #14 #15
E12.5 E14.5 E12.5 E13.5 E13.5 E14.5
Sampling Day 1 Sampling Day 2 Sampling Day 3
F
Quality control (on seperate samples)
Preprocessing (on combined dataset)
Seurat scaling
Seurat log-transformation
Seurat cell cycle scoring
1st level (Major Cell Types)
Feature selection
Seurat regression
Seurat PCA
Hierarchical clustering
Seurat marker genes
Transfer data to Python
Scanpy PCA
BBKNN correction
Scanpy UMAP
Visualization
2nd level (Fibroblasts)
Feature selection
Seurat regression
Transfer data to Python
Scanpy PCA
BBKNN correction
Scanpy UMAP
Scanpy Leiden clustering
scVelo velocity analysis
Visualization
Transfer clustering to R
Seurat marker genes
2nd level (Epidermis)
Feature selection
Seurat regression
Seurat PCA
Seurat tSNE
Hierarchical clustering
Seurat marker genes
Transfer data to Python
scVelo velocity analysis
CellRank lineage drivers
Visualization
Receptor-Ligand-Analysis
Custom pipeline (see Fig. S5)
G E12.5
H E13.5
I E14.5
M

**Figure S1. Details of experimental approach and quality control, Related to Figure 1**

(A) Overview of the experimental workflow.
(B) Sampling area highlighted in blue.
(C) Representative photos and H&E sagital sections of embryos at the analysed embryonic time points.
(D) Sample contributions (n = 5 biological replicates) per timepoint overlayed on combined UMAP.
(E) Violin plots showing a uniform distribution of QC measures (reads/cell, genes/cell and percentage of mitochondrial reads among all reads) among all 15 biological replicates (upper 3 panels). Violin plots showcasing the approach of retrospective sex determination (lower 2 panels). Only male embryos show reads from Y chromosome.
(F) Flow diagram summarizing the analysis workflow. Black font marks steps that were performed in R and blue font marks steps that were performed in Python.
(G-I) H&E stained tissue sections of dorsal skin. Zoom in on epidermis (includind a developing hair follicle in (H)). M marks developing muscle layers (for orientation in the tissue). Scale bars, 100µm.

#### Figure S2

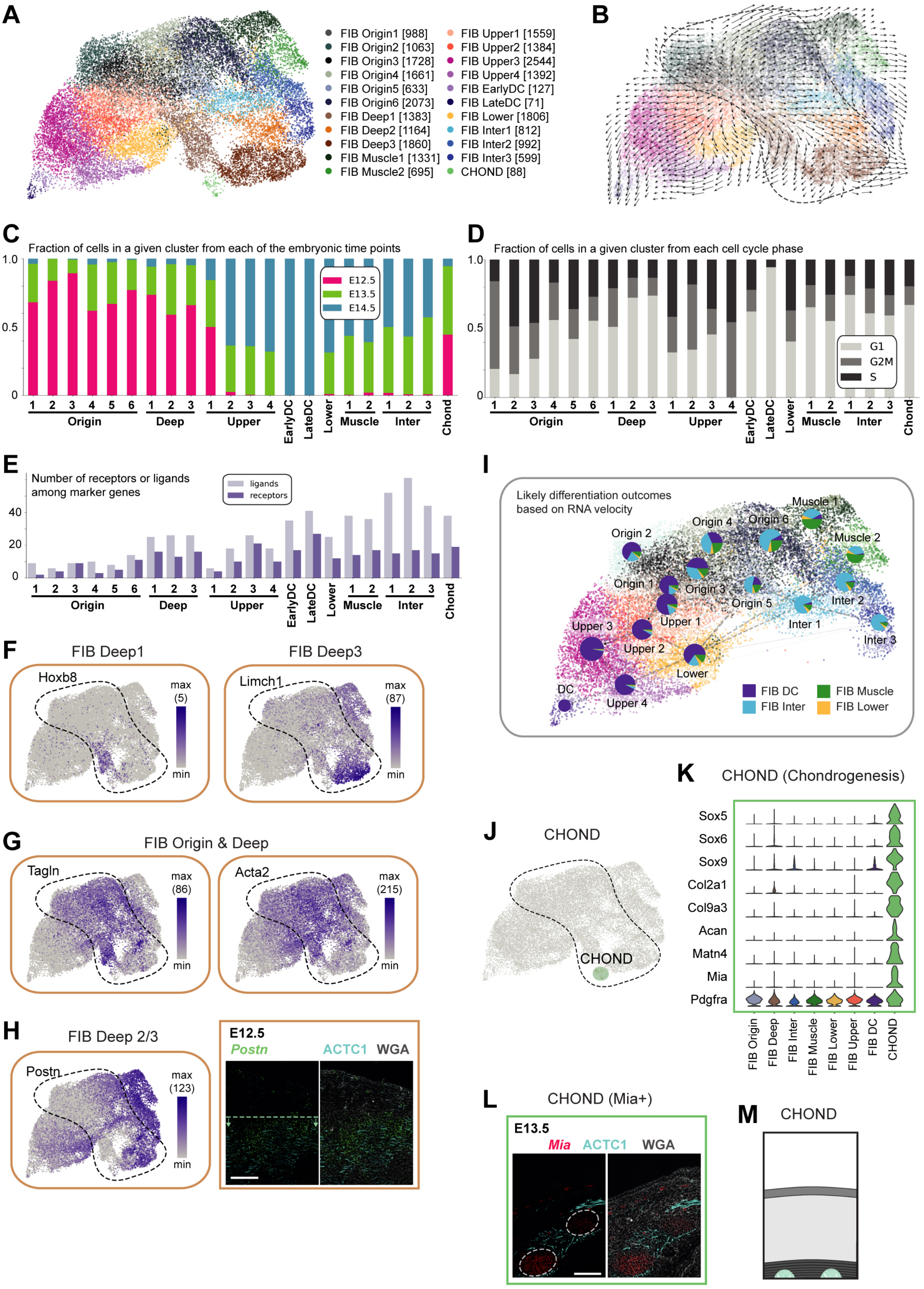
A
FIB Origin1 [988]
FIB Origin2 [1063]
FIB Origin3 [1728]
FIB Origin4 [1661]
FIB Origin5 [633]
FIB Origin6 [2073]
FIB Deep1 [1383]
FIB Deep2 [1164]
FIB Deep3 [1860]
FIB Muscle1 [1331]
FIB Muscle2 [695]
FIB Upper1 [1559]
FIB Upper2 [1384]
FIB Upper3 [2544]
FIB Upper4 [1392]
FIB EarlyDC [127]
FIB LateDC [71]
FIB Lower [1806]
FIB Inter1 [812]
FIB Inter2 [992]
FIB Inter3 [599]
CHOND [88]
B
C
Fraction of cells in a given cluster from each of the embryonic time points
E12.5
E13.5
E14.5
Origin
Deep
Upper
EarlyDC
LateDC
Lower
Muscle
Inter
Chond
D
Fraction of cells in a given cluster from each cell cycle phase
G1
G2M
S
E
Number of receptors or ligands among marker genes
ligands
receptors
F
FIB Deep1
Hoxb8
FIB Deep3
Limch1
max
min
G
FIB Origin & Deep
Tagln
Acta2
H
FIB Deep 2/3
Postn
E12.5
Postn
ACTC1
WGA
I
Likely differentiation outcomes based on RNA velocity
Muscle 1
Origin 4
Origin 6
Origin 2
Muscle 2
Origin 1
Origin 3
Origin 5
Inter 2
Upper 3
Upper 1
Inter 1
Inter 3
Upper 2
Lower
DC
Upper 4
FIB DC
FIB Inter
FIB Muscle
FIB Lower
J
CHOND
K
CHOND (Chondrogenesis)
Sox5
Sox6
Sox9
Col2a1
Col9a3
Acan
Matn4
Mia
Pdgfra
FIB Origin
FIB Deep
FIB Inter
FIB Muscle
FIB Lower
FIB Upper
FIB DC
CHOND
L
CHOND (Mia+)
E13.5
Mia
ACTC1
WGA
M
CHOND

**Figure S2. Deconstruction of fibroblast heterogeneity at E12.5 (expression and location), Related to Figure 2**

(A) UMAP visualization of fibroblasts, colored according to fibroblast subclustering. Cell numbers per cluster are displayed in square brackets.
(B) UMAP, colored according to fibroblast subclustering and overlayed with RNA velocity vectors that predict developmental trajectories.
(C) Bar plot visualizing the contribution of each embryonic time point to each fibroblast subcluster.
(D) Bar plot visualizing the contribution of each (predicted) cell cycle phase to each fibroblast subcluster.
(E) Bar plot showing the number of ligands and receptors among the marker genes of each fibroblast subcluster.
(F) Expression pattern of additional *FIB Deep* marker genes projected onto UMAP. Maximum number of mRNA copies detected per cell is presented in brackets to provide an idea of the absolute abundance of the marker gene.
(G) Expression pattern of additional marker genes shared between *FIB Origin* and *FIB Deep* projected onto UMAP. Maximum number of mRNA copies detected per cell is presented in brackets to provide an idea of the absolute abundance of the marker gene.
(H) Expression pattern of *Postn* projected onto UMAP (left panel). *Postn* mRNA staining plus ACTC1 protein staining (right panel) in dorsal E12.5 skin. Dashed lines with arrows highlight the region with highest expression. Counterstained with WGA. Microscope images originate from larger tile scan (n = 3 mice). Scale bars, 100µm.
(I) Directed PAGA plot showing the most likely differentiation outcomes of each fibroblast subpopulation based on RNA velocity. Cells are colored according to fibroblast subclustering and pie chart segments are colored according to the differentiation outcomes.
(J) *FIB CHOND* cells highlighted on the UMAP.
(K) Violin Plots showing expression of *FIB CHOND* marker genes.
(L) *Mia* mRNA and *ACTC1* protein staining in dorsal E13.5 skin reveals location of *FIB CHOND* cells. Counterstained with WGA. Microscope image originates from larger tile scan (n = 3 mice). Scale bar, 100µm.
(M) Scheme summarizing the location of *FIB CHOND* cells within the tissue at E12.5.

### Figure S3

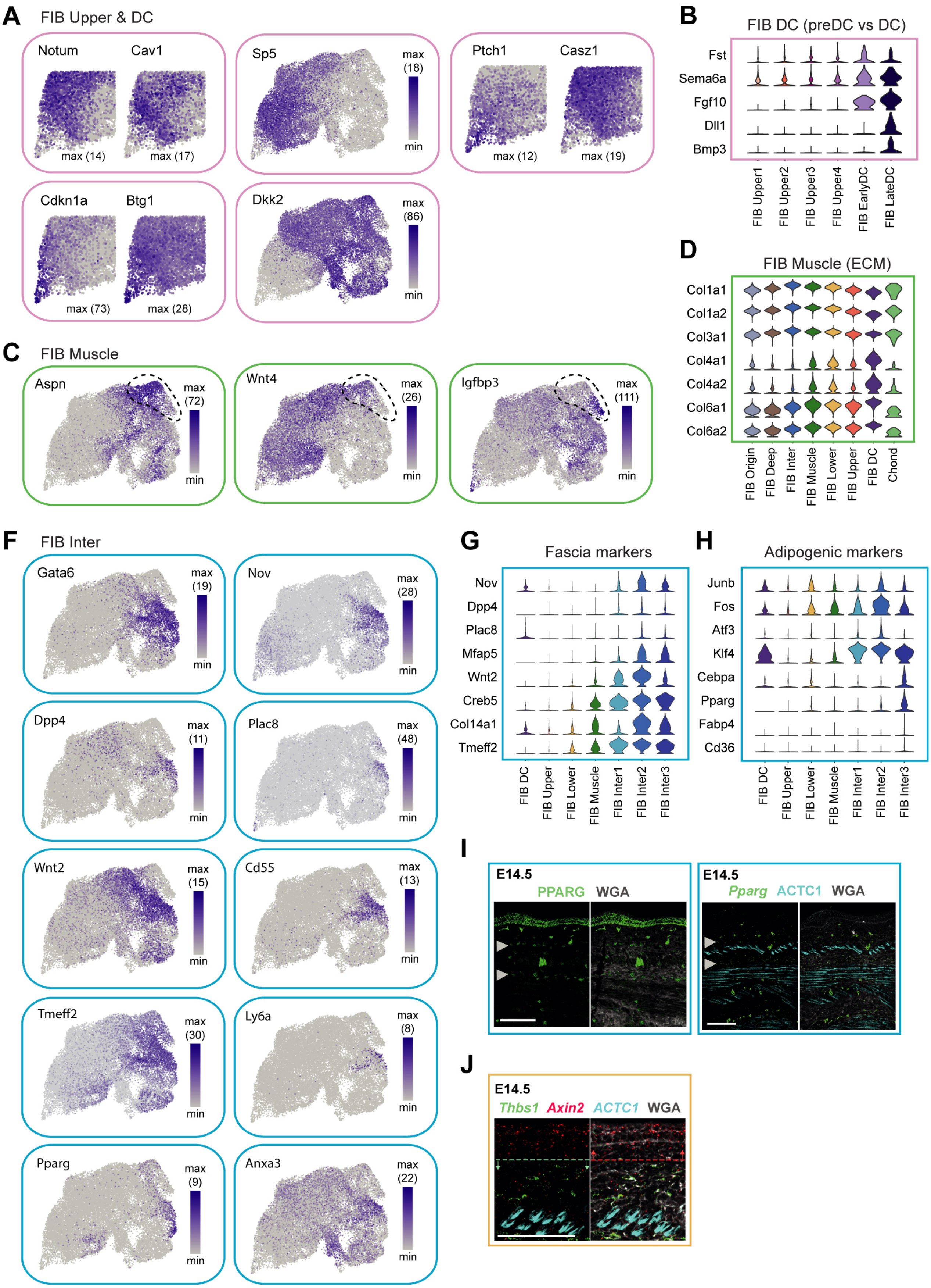
A
FIB Upper & DC
Notum
Cav1
max (14)
max (17)
Sp5
max
(18)
min
Ptch1
Casz1
max (12)
max (19)
Cdkn1a
Btg1
max (73)
max (28)
Dkk2
max
(86)
min
B
FIB DC (preDC vs DC)
Fst
Sema6a
Fgf10
Dll1
Bmp3
FIB Upper1
FIB Upper2
FIB Upper3
FIB Upper4
FIB EarlyDC
FIB LateDC
C
FIB Muscle
Aspn
max
(72)
min
Wnt4
max
(26)
min
Igfbp3
max
(111)
min
D
FIB Muscle (ECM)
Col1a1
Col1a2
Col3a1
Col4a1
Col4a2
Col6a1
Col6a2
FIB Origin
FIB Deep
FIB Inter
FIB Muscle
FIB Lower
FIB Upper
FIB DC
Chond
F
FIB Inter
Gata6
max
(19)
min
Nov
max
(28)
min
Dpp4
max
(11)
min
Plac8
max
(48)
min
Wnt2
max
(15)
min
Cd55
max
(13)
min
Tmeff2
max
(30)
min
Ly6a
max
(8)
min
Pparg
max
(9)
min
Anxa3
max
(22)
min
G
Fascia markers
Nov
Dpp4
Plac8
Mfap5
Wnt2
Creb5
Col14a1
Tmeff2
FIB DC
FIB Upper
FIB Lower
FIB Muscle
FIB Inter1
FIB Inter2
FIB Inter3
H
Adipogenic markers
Junb
Fos
Atf3
Klf4
Cebpa
Pparg
Fabp4
Cd36
FIB DC
FIB Upper
FIB Lower
FIB Muscle
FIB Inter1
FIB Inter2
FIB Inter3
I
E14.5
PPARG WGA
E14.5
Pparg ACTC1 WGA
J
E14.5
Thbs1 Axin2 ACTC1 WGA

**Figure S3. Deconstruction of fibroblast heterogeneity at E13.5 and E14.5 (expression and location), Related to Figure 3**

(A) Expression pattern of additional *FIB Upper / DC* marker genes projected onto UMAP.
(B) Violin Plots showing expression of DC marker genes.
(C) Expression pattern of additional *FIB Muscle* marker genes projected onto UMAP.
(D) Violin Plots showing expression of ECM-related genes.
(F) Expression pattern of additional *FIB Inter* marker genes projected onto UMAP.
(G) Violin Plots showing expression of fascia-related marker genes.
(H) Violin Plots showing expression of adipogenesis-related marker genes.
(I) PPARG protein staining (left panel) and *Pparg* mRNA staining plus ACTC1 protein staining (right panel) in dorsal E14.5 skin. Arrowheads mark layers with maximal staining. Counterstained with WGA. Microscope images originate from larger tile scan (n = 3 mice). Scale bars, 100µm.
(J) *Thbs1* and *Axin2* mRNA staining plus ACTC1 protein staining in dorsal E14.5 skin. Arrowheads mark layers with maximal staining. Counterstained with WGA. Dashed lines with arrows highlight the region with highest expression. Microscope images originate from larger tile scan (n = 3 mice). Scale bars, 100µm.

### Figure S4

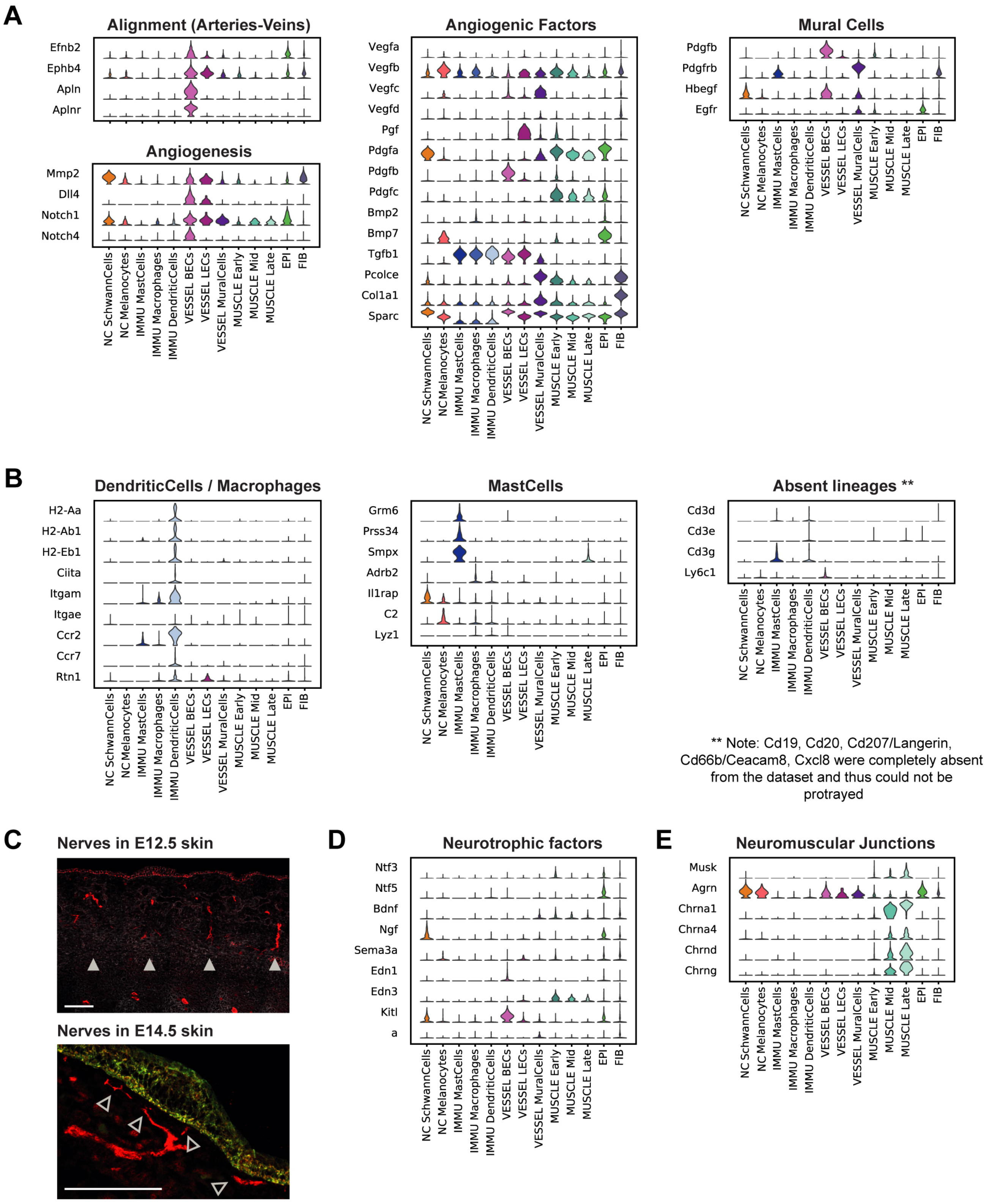
A
Alignment (Arteries-Veins)
Efnb2
Ephb4
Apln
Aplnr
Angiogenesis
Mmp2
Dll4
Notch1
Notch4
Angiogenic Factors
Vegfa
Vegfb
Vegfc
Vegfd
Pgf
Pdgfa
Pdgfb
Pdgfc
Bmp2
Bmp7
Tgfb1
Pcolce
Col1a1
Sparc
Mural Cells
Pdgfb
Pdgfrb
Hbegf
Egfr
NC SchwannCells
NC Melanocytes
IMMU MastCells
IMMU Macrophages
IMMU DendriticCells
VESSEL BECs
VESSEL LECs
VESSEL MuralCells
MUSCLE Early
MUSCLE Mid
MUSCLE Late
EPI
FIB
B
DendriticCells / Macrophages
H2-Aa
H2-Ab1
H2-Eb1
Ciita
Itgam
Itgae
Ccr2
Ccr7
Rtn1
MastCells
Grm6
Prss34
Smpx
Adrb2
Il1rap
C2
Lyz1
Absent lineages **
Cd3d
Cd3e
Cd3g
Ly6c1
** Note: Cd19, Cd20, Cd207/Langerin, Cd66b/Ceacam8, Cxcl8 were completely absent from the dataset and thus could not be protrayed
C
Nerves in E12.5 skin
Nerves in E14.5 skin
D
Neurotrophic factors
Ntf3
Ntf5
Bdnf
Ngf
Sema3a
Edn1
Edn3
Kitl
a
E
Neuromuscular Junctions
Musk
Agrn
Chrna1
Chrna4
Chrnd
Chrng

**Figure S4. Cell types contributing to embryonic skin besides fibroblasts and keratinocytes, Related to Figure 5**

(A) Violin Plots showing expression of markers used to further characterize the captured vessel-associated cells.
(B) Violin Plots showing expression of markers used to further characterize the captured immune cells.
(C) PPARG and KRT5 protein staining in dorsal E12.5 and E14.5 skin, respectively. Filled arrowheads mark thick nerve bundles traversing dermis at the height of each vertebrae (asterisks). Empty arrowheads mark nerve endings extending towards epidermis. Counterstained with WGA. Microscope images originate from larger tile scans (n = 3 mice). Scale bars, 100µm.
(D) Violin Plots showing expression of markers used to further characterize the captured neural crest-derived cells.
(E) Violin Plots showing expression of markers used to further characterize the captured muscle cells.

#### Figure S5

##### Identification

(1) Identify marker genes for each cluster
(2) Identify receptors and ligands among those marker genes (by querying rec-lig-database)
(3) Find potential rec-lig-pairs among expressed receptors and ligands (by querying rec-lig-database)

↓

##### Filtering

(4) Run statistical test to determine how likely it is to find the obseverd amount of rec-lig-pairs by chance
(5) Keep rec-lig-pairs with $p \leq 0.05$ (Benjamini-Hochberg-corrected p-value)
(6) Combine all significant rec-lig-pairs into Table S3
(7) Add functional annotations for each of the included receptors and ligands
(8) Assign scores to each rec-lig-pair for their potential involvement in vascular development, neuronal development, immune cell development / recruitment (Has any functional link been established in the literature? Does the expression pattern match?)
(9) Keep pairs with score ≥ 0.75 for any of the aforementioned processes

↓

##### Visualization

(10) Combine all filtered rec-lig-pairs into the central panel of Figure 5A-C
(11) Plot expression of all included receptors and ligands in left and right panel of Figure 5A-C (heatmaps)
(12) A subset of genes from Figure 5A-C with high expression in either dermal condensate or hair placode was selected manually for display in Figure S6K

**Figure S5. Workflow for receptor-ligand analysis, Related to Figure 6**

Flow diagram summarizing the analysis workflow that was followed to obtain the presented receptor-ligand pairs in **Figure 5** as well as **Table S3**.

### Figure S6

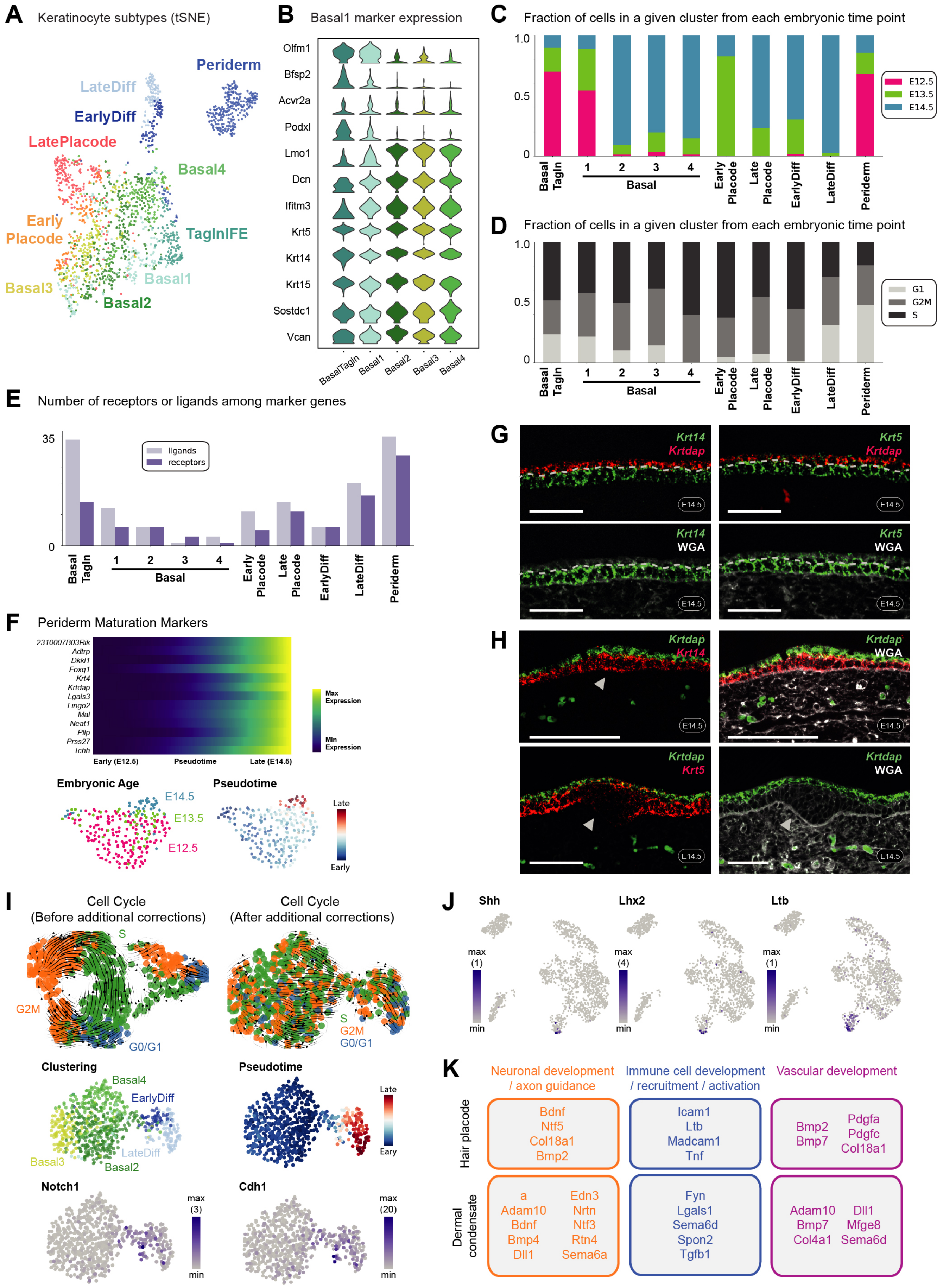
A
Keratinocyte subtypes (tSNE)
Periderm
LateDiff
EarlyDiff
LatePlacode
Basal4
Early Placode
TaglnIFE
Basal1
Basal3
Basal2
B
Basal1 marker expression
Olfm1
Bfsp2
Acvr2a
Podxl
Lmo1
Dcn
Ifitm3
Krt5
Krt14
Krt15
Sostdc1
Vcan
BasalTagln
Basal1
Basal2
Basal3
Basal4
C
Fraction of cells in a given cluster from each embryonic time point
1.0
0.5
0
E12.5
E13.5
E14.5
Basal Tagln
1
2
3
4
Basal
Early Placode
Late Placode
EarlyDiff
LateDiff
Periderm
D
Fraction of cells in a given cluster from each embryonic time point
G1
G2M
S
E
Number of receptors or ligands among marker genes
35
0
ligands
receptors
F
Periderm Maturation Markers
2310007B03Rik
Adtrp
Dkkl1
Foxq1
Krt4
Krtdap
Lgals3
Lingo2
Mal
Neat1
Pllp
Prss27
Tchh
Max Expression
Min Expression
Early (E12.5)
Pseudotime
Late (E14.5)
Embryonic Age
Pseudotime
E14.5
E13.5
E12.5
Late
Early
G
Krt14
Krtdap
Krt5
Krtdap
Krt14
WGA
Krt5
WGA
E14.5
H
Krtdap
Krt14
Krtdap
WGA
Krtdap
Krt5
Krtdap
WGA
I
Cell Cycle
(Before additional corrections)
Cell Cycle
(After additional corrections)
S
G2M
G0/G1
Clustering
Basal4
EarlyDiff
LateDiff
Basal3
Basal2
Pseudotime
Late
Eary
Notch1
max
(3)
min
Cdh1
max
(20)
min
J
Shh
Lhx2
Ltb
max
(1)
min
max
(4)
min
K
Neuronal development / axon guidance
Immune cell development / recruitment / activation
Vascular development
Hair placode
Bdnf
Ntf5
Col18a1
Bmp2
Icam1
Ltb
Madcam1
Tnf
Bmp2
Bmp7
Pdgfa
Pdgfc
Col18a1
Dermal condensate
a
Adam10
Bdnf
Bmp4
Dll1
Edn3
Nrtn
Ntf3
Rtn4
Sema6a
Fyn
Lgals1
Sema6d
Spon2
Tgfb1
Adam10
Bmp7
Col4a1
Dll1
Mfge8
Sema6d

**Figure S6. Epidermal development from a single basal layer towards a HF-inducing and stratified epithelium, Related to Figure 7**

(A) tSNE visualization of all keratinocytes, colored according to keratinocyte subclustering.
(B) Violin Plot showing overlapping signatures in *EPI Basal1* population.
(C) Bar plot visualizing the contribution of each embryonic time point to each keratinocyte subcluster.
(D) Bar plot visualizing the contribution of each (predicted) cell cycle phase to each keratinocyte subcluster.
(E) Bar plot showing the number of ligands and receptors among the marker genes of each keratinocyte subcluster.
(F) Heatmap of genes related to periderm maturation (upper panel). A pseudotime was modeled based on *EPI Periderm* cells from all embryonic time points and pseudotime-dependent genes were determined. Genes peaking latest in pseudotime were chosen for display on heatmap. UMAP of periderm cells, colored according to embryonic age (lower left panel) or pseudotime (lower right panel), respectively.
(G) *Krt14* and *Krtdap* mRNA staining (left panels) and *Krt5* and *Krtdap* mRNA staining (right panels) in dorsal E14.5 skin reveals differing expression pattern of Krt5 and Krt14 in suprabasal layer. Dashed line outlines basal-suprabasal border. Counterstained with WGA. Microscope images originate from larger tile scans (n = 3 mice). Scale bars, 50µm.
(H) *Krt14*, *Krt5* and *Krtdap* mRNA staining in dorsal E14.5 skin confirms downregulation of *Krt5* and maintenance of *Krt14* in the developing hair placode (arrowhead) and showcases a continous layer of suprabasal *Krtdap*$^{+}$ cells covering the hair placode. Counterstained with WGA. Microscope image originates from larger tile scan (n = 3 mice). Scale bars, 50µm.
(I) UMAP visualization of keratinocyte subset (*EPI Basal1-4*, *EPI EarlyDiff*, and *EPI LateDiff* cells from E14.5) used to delineate epidermal stratification. Row one: UMAP before and after additional corrections for cell cycle (see **Methods**), colored according to (predicted) cell cycle phase and overlayed with velocity vectors predicting developmental trajectories. Row two: UMAP (from upper right panel), colored according to keratinocyte subclustering (left panel) or pseudotime (right panel), respectively. Row three: Expression of *Notch1* and *Cdh1* projected onto the UMAP from the upper right panel. Maximum number of mRNA copies detected per cell is presented in brackets to provide an idea of the absolute abundance of the marker gene.
(J) Expression of placode markers projected onto the UMAP from **Figure 6A**.
(K) Summary scheme with selected hair placode- and dermal condensate-derived factors promoting neuoronal, immune cell, and vascular development.
